# Heterozygous RELA mutations cause early-onset systemic lupus erythematosus by hijacking the NF-κB pathway towards transcriptional activation of type-I Interferon genes

**DOI:** 10.1101/2020.04.27.046102

**Authors:** Laura Barnabei, Hicham Lamrini, Mathieu Castela, Nadia Jeremiah, Marie-Claude Stolzenberg, Loïc Chentout, Sidonie Jacques, Amine Bouafia, Aude Magérus-Chatinet, Matthieu Moncan, Sajedeh Mirshahvalad, Olivier Pellé, Vincent Bondet, Darragh Duffy, Mélanie Parisot, Marc Bras, Carolina Uggenti, Rémi Salomon, Christine Bodemer, Marion Rabant, Marina Cavazzana, J.J. Miner, Alexandre Belot, Miguel Hié, Capucine Picard, Brigitte Bader-Meunier, Sven Kracker, Frédéric Rieux-Laucat

## Abstract

Systemic Lupus Erythematosus (SLE) is an autoimmune and inflammatory disease characterized by uncontrolled production of autoantibodies and inflammatory cytokines such as the type-I interferons. Due to the lack of precise pathophysiological mechanisms, treatments are based on broad unspecific immunossupression. To identify genetic factors associated with SLE we performed whole exome sequencing and identified two *RELA* heterozygous activating mutations in 3 early-onset and familial SLE cases. The corresponding RELA/p65 mutant were abundant in the nucleus but poorly activate transcription of genes controlled by NF-κB consensus sequences. The co-expression of the mutant and wild-type RELA/p65 strongly activated the expression of genes controlled by the IFNα-consensus sequences. These molecular mechanisms lead to the overproduction of type-I IFN in the patients’ cells. Our findings highlight a novel mechanism of autoimmunity where these new RELA mutants are transactivating the type-I IFN genes and are thus promoting type-I interferon production and early-onset SLE, thereby paving the way to the identification of new and specific therapeutic targets.

**Summary:** Heterozygous RELA mutations are associated with Systemic Lupus Erythematosus, with increased expression of genes controlled by the IFNα-consensus sequences.

## Introduction

Systemic Lupus Erythematosus (SLE) is a systemic multiorgan autoimmune and inflammatory disease that can affect multiple organs, with a near-universal pathophysiological mechanism involving a breakage in B cell tolerance leading to uncontrolled production of auto-antibodies against nuclear elements(Tsokos et al. 2016). Defective clearance of apoptotic bodies has been depicted as a possible trigger through exacerbated exposition of nuclear antigens (DNA and chromatin associated proteins) to B cells. This is illustrated by the high incidence (>80%) of SLE onset in patients with complete C1q deficiency and to a lesser extent in patients with other complement deficiencies(Macedo and Isaac 2016). Besides the stimulation of autoreactive B cells, the exposition of nuclear antigens, notably of nucleic acids, can stimulate the Toll-like receptors (TLRs), thereby triggering an inflammatory response that can contribute to the breakage of tolerance. One of the central players appears to be the type-I interferon (IFN-I) family, principally IFN-alpha (IFN-α) and IFN-beta (IFN-β)(Crow 2014). Elevated levels of IFN-I in the blood of SLE patients has been long recognized(Hooks et al. 1979). The subsequent identification of Interferon-Stimulated Genes (ISGs) led to the demonstration of a broad ISG transcript signature in Peripheral Blood Mononuclear Cells (PBMCs) from SLE patients(Baechler et al. 2003; Bennett et al. 2003; Crow, Kirou, and Wohlgemuth 2003; Han et al. 2003). Whereas several hundred gene transcripts define this signature, most are highly correlated, and assessment of as few as 5 transcripts quantified by real-time PCR is now used as a biomarker reflecting the functional effect of IFN-I *in vivo* in SLE patients(Yao et al. 2009). The accuracy of the ISG signature was further confirmed using the recent development of digital ELISA, allowing a very sensitive measurement of IFN-I molecules in the sera of patients and showing a very good correlation between elevated ISGs and IFN-I concentrations(Rodero et al. 2017). The central role of IFN-I in the pathogenesis of lupus was also highlighted by increased incidence of SLE or SLE-like onset in rare monogenic diseases with gene defects driving IFN-I overproduction defined as Type I interferonopathies(Lee-Kirsch 2017). Aicardi-Goutières Syndrome (AGS) is the prototype of these interferonopathies and is caused by recessive or dominant mutations of genes involved in intracellular nucleic acids homeostasis (such as *TREX1, SAMHD1, RNASEH2A/B/C* and *ADAR1*), or in nucleotide sensing such as *IFIH1* /MDA5(Crow and Manel 2015; Roers, Hiller, and Hornung 2016; Lee-Kirsch 2017). Interestingly, activating mutations in *TMEM173*/STING, an adaptor of the DNA sensor cGAS, also lead to IFN-I overproduction and are sometimes associated with chilblain lupus or SLE-like features(Jeremiah et al. 2014; Liu et al. 2014; Konig et al. 2017). Control of extracellular DNA is also important as *DNASE1L3* mutations have been found in a familial form of SLE(Al-Mayouf et al. 2011) as well as in the lupus-prone MRL and NZB/W F1 mice(Wilber et al. 2003). Disturbance of extra or intra cellular nucleic acid homeostasis converges to IRF3 and NF-κB activation, resulting in expression of inflammatory genes including the IFN-I family(Tsokos et al. 2016). Among the numerous risk loci associated with SLE(Hom et al. 2008; International Consortium for Systemic Lupus Erythematosus et al. 2008; Gateva et al. 2009), common genetic variants, affecting genes involved in the control of the NF-κB pathway (*TNFAIP3*/A20)(Graham et al. 2008), have been reported in genome-wide association (GWAS) studies of SLE cohorts. Moreover, SLE has been described in one patient with a heterozygous loss-of-function mutation of the *TNFAIP3* gene, leading to A20 haploinsufficiency (HA20)(Aeschlimann et al. 2018). Finally, the *TNFAIP3* gene is located nearby the HLA locus, one of the first genetic loci linked to autoimmunity, in particular to SLE(Graham et al. 2007). These results suggest that a dysregulation of the NF-κB pathway could be involved in the pathogenesis of some SLE patients.

The present study supports an initial role for dysregulation of the NF-κB pathway and type-I IFN overproduction in the pathogenesis of SLE. Heterozygous *RELA* mutations cause the expression of mutant RELA/p65 proteins responsible for hijacking of the NF-κB activity towards the activation of inflammatory cytokine genes.

## Results

### Clinical and biological presentation

The index patient from family 1 (P1) is a female born to an Algerian couple originating from the same Algerian village but without known or identified consanguinity. At 1.5 years of age, she was diagnosed as having SLE, with malar rash, oral ulcers, polyarthralgia, alopecia and fever. Blood analyses evidenced low C4 values, increased levels of anti–double-stranded DNA, anti-SSa, and anti-SSb antibodies and rheumatoid factor (Table. S1). At 12 years of age, she developed severe cutaneous vasculitis of toes and fingers (Fig. S1). All attempts at decreasing corticosteroids were associated with severe cutaneous flares and fever, despite the association of mycophenolate mofetyl or rituximab. At 18 years, corticosteroids were still required. The patient developed a prolonged severe hypogammaglobulinemia and B cell lymphopenia after anti-CD20 antibody infusion. Before treatment the patient’s globulin level was elevated (IgG: 19 g/L, IgA: 2.26 g/L and IgM: 2.53 g/L). The IgG levels dropped to 5 g/L and IgM were undetectable 15 months and 3 years after initiation of treatment. Of note, the IgA level remained within normal values (Table. S2). CD19^+^ B cells became undetectable one month after the first infusion of anti-CD20 antibody and reconstitution of the B cell pool was still not observed 3 years later. Despite subcutaneous immunoglobulins replacement, the patient suffered from chronic suppurative otitis due to *Hemophilus Influenzae* infections. The P1’s mother (P2) developed SLE at 33 years of age, and presented with severe non-erosive polyarthritis, oral ulcers, pericarditis and class IV lupus-nephritis associated with positive antinuclear antibodies (ANA) and anti-double-stranded DNA (Table S.1). At 48 years of age, no cutaneous involvement had occurred.

The patient from the second family (P3) is a boy born to non-consanguineous parents of French Caucasian ethnicity. Both parents are healthy. He presented at 9 years of age with a lupus rash, oral and skin ulcers on limbs, arthralgia, positive ANA and anti-double-stranded DNA. SLE was diagnosed and hydroxycholoroquine in combination with a low-dose prednisone (10 mg/d) was started. He further developed Class IV lupus nephritis and pericarditis. At age 17 years, the patient was in clinical and biological remission with mycophenolate mofetil given orally.

### Identification of heterozygous *RELA* variants in 3 SLE patients

Whole exome sequencing (WES) of DNA from total blood samples was performed in both pediatric patients and their parents. Rare deleterious (predicted by the Sift and Polyphen software and a CADD score >25) heterozygous variations in *RELA* were detected by filtering WES data for autosomal dominant (family 1) and *de novo* variations (family 2). These variants were not previously described neither in our in-house (>12 000 WES) nor in the genome aggregated databases (gnomAd: 123,136 exomes and 15,496 genomes from unrelated individuals sequenced as part of various disease-specific and population genetic studies). In the DNA from P1 and P2, a heterozygous C to A variation at position 256 (GRCh37; NM_021975.3 c.256 C>A) in exon 4 of the *RELA* gene was detected (Fig. 1A). The variation encodes a histidine to asparagine substitution at position 86 (p.H86N) located in the REL-Homoly-Domain (RHD) (Fig. 1B). This histidine residue within the RHD of RELA/p65 is evolutionarily highly conserved (Fig.S2). A heterozygous C to T nonsense variation at position 985 (GRCh37; NM_021975.3 c.985 C>T) in exon 10 of the *RELA* gene was detected in P3 (Fig. 1A). The variation leads to a truncated protein, substituting an arginine to an early STOP at the position 329 (p.R329X). Sanger sequencing confirmed the mutation in the patient’s cDNA, which was not present in both parents (Fig. 1A). Furthermore, both mutations were detected at the genomic DNA level (Fig. S3).

**Figure 1.**
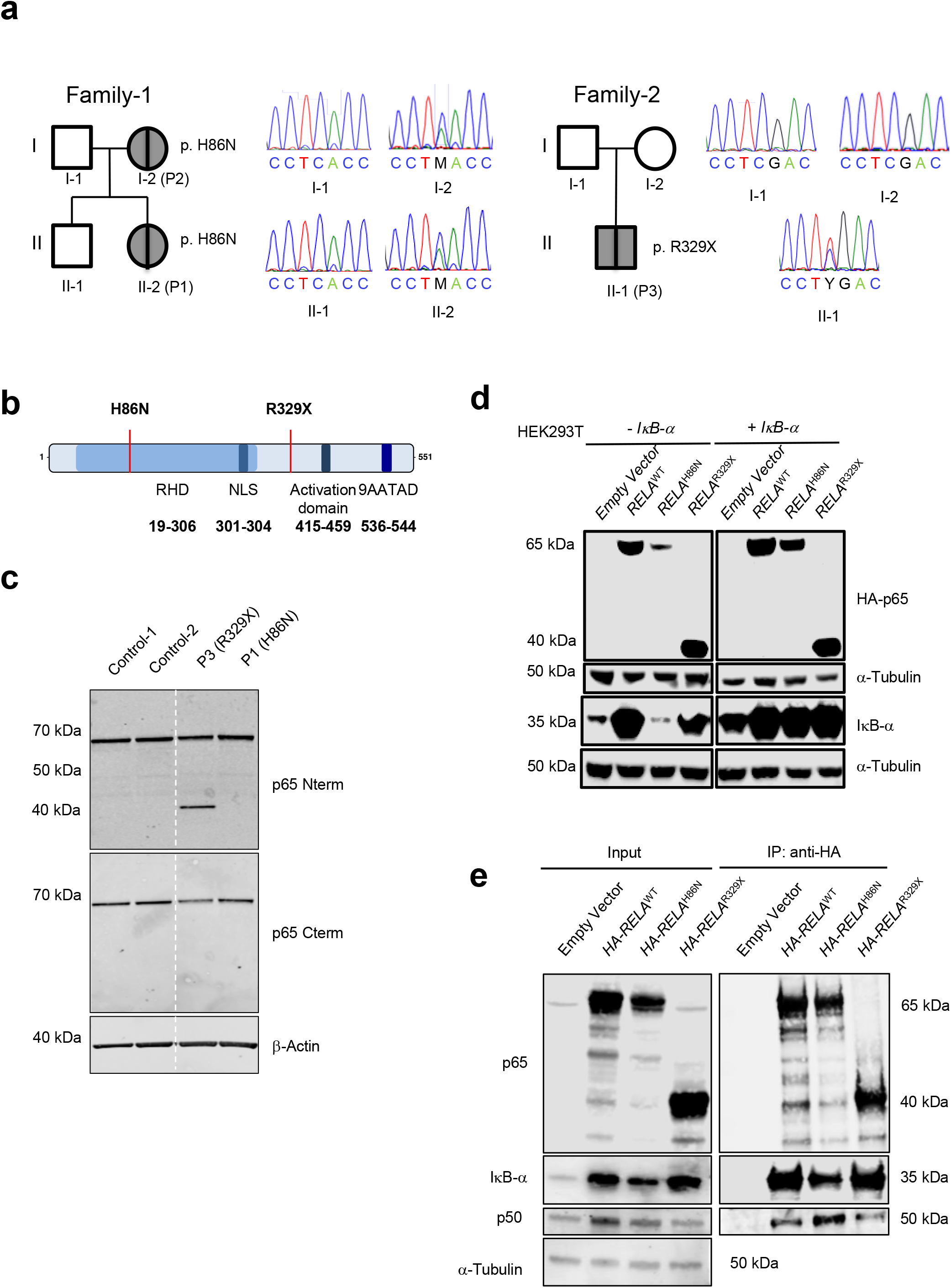
Heterozygous RELA mutations segregated with Systemic Lupus Erythematosus and interact with IκB-α and NF-κB1/p50. (**a**) Pedigree and Sanger sequencing of cDNA of the family-1 and 2. In family 1 a heterozygous RELA c.256 C>A missense mutation predicted to result in the p.H86N mutated RELA/p65 protein was found in P1 and P2. In family 2 a heterozygous RELA c.985 C>T nonsense mutation predicted to result in the p.R329X truncated RELA/p65 protein was found in P3. (**b**) RELA/p65 protein structure with indicated mutations positions. RHD = REL-Homology-Domain; NLS = Nuclear localization signal; 9AATAD = 9 amino acid transactivation domain. (**c**) Expression of RELA/p65 in activated T-cell blasts of probands and healthy relative controls by Western Blot analysis using antibodies specific for N-terminal (Nterm) and C-terminal (Cterm) domains of RELA/p65. Anti-β-Actin was used as an internal control. (**d**) Immuno-blotting of HEK293T cell lysates co-transfected with constructs expressing N-terminal HA-tagged RELA/p65 and IκB-α with a ratio of 3:1. Anti-HA-RELA/p65, anti-IκB-α and anti-α-tubulin antibodies were used to assess protein expression. (**e**) Immunoblotting of cell lysates from co-transfected HEK293T cells with an IκB-α construct and HA-RELA/p65 constructs performed as indicated. Proteins were immuno-precipitated with anti-HA antibody. Initial lysates and immune-precipitated proteins were subjected to immunoblotting with anti-RELA/p65, anti-IκB-α anti-NF-κB1/p50 and anti-@-tubulin antibodies. A representative experiment is depicted.

To analyze the impact of *RELA* variations on RELA/p65 protein expression, immunoblotting was performed on protein cell lysates from patients’ activated T-cell blasts using RELA/p65-specific antibodies (Fig. 1C). The RELA/p65 protein abundance was only slightly decreased for P1 carrying the *RELA*^H86N^ allele when compared to controls suggesting the presence of a *RELA*^H86N^ protein (Fig. 1C). An approximate 40-42 kDa RELA/p65 protein together with a ~50% reduction in RELA/p65 Cterm wild-type protein expression compared to controls was detected in cell extracts from P3 carrying the RELA/p65^R329X^ allele. This truncated RELA/p65 protein was not present in controls (Fig. 1C and Fig. S4).

Together, these data indicate that the heterozygous variants in *RELA* in the SLE patients, result in the expression of RELA/p65^H86N^ and RELA/p65^R329X^ proteins affecting the RHD or leading to a truncated RELA/p65 lacking most of the transactivation domain, respectively.

### Increased nuclear localization of RELA/p65 mutants, with DNA binding capacity, in lymphocytes from SLE patients

Next, we investigated the intracellular localization of RELA/p65 proteins. To distinguish between truncated, that miss the C-terminal part, and full-length RELA/p65 proteins we performed staining with antibodies recognizing either the N-terminal or C-terminal domains of RELA/p65 (Fig 2A). Unstimulated activated T-cell blasts from P1 (RELA/p65^H86N^) showed increased nuclear staining using both antibodies, whereas unstimulated activated T-cell blasts from P3 (RELA/p65^R329X^) displayed increased nuclear staining with the antibody recognizing the N-terminal domain of RELA/p65 (Fig. 2A). Both patients’ activated T-cell blasts showed increased nuclear staining of N- and C-terminal domains of RELA/p65 compared to controls when stimulated for 30 min with TNF-α or PMA-ionomycin (Fig. 2A and Fig. S5).

**Figure 2.**
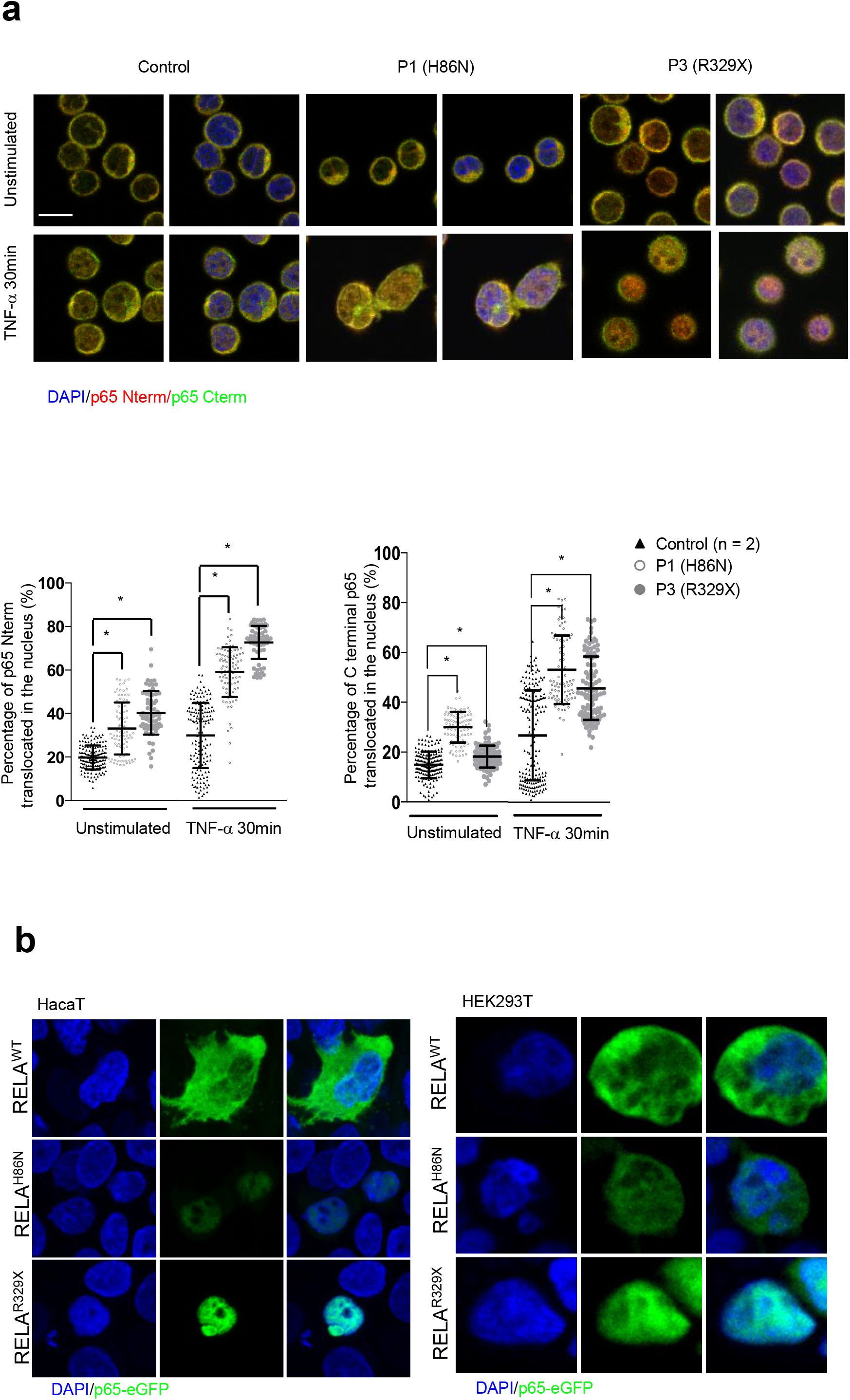
RELA/p65 proteins are located in both cytoplasmic and nuclear compartments. (**a**) Immunofluorescence of activated T-cell blasts from P1 (H86N), P3 (R329X) and healthy donor stimulated 30 minutes with TNF-α and analyzed by confocal microscopy using N-terminus (Nterm)-specific anti-RELA/p65 (Red), C-terminus (Cterm)-specific anti-RELA/p65 (Green) antibodies, and DAPI (4’-6-diamidino-2-phenylindole). Top panels: Representative pictures. Bottom Panel: Quantification of the RELA/p65 nuclear localization with the N-terminal-specific antibody and C-terminal-specific antibody. Mann-Whitney Wilcoxon, \**P*<0.05; mean±SD. (**b**) Immunofluoresence of HacaT and HEK293T cells transfected with the different RELA constructs analyzed by confocal microscopy. Scale bar = 5®m.

We then transfected HacaT or HEK293T cells with the wild-type, H86N or R329X mutated RELA/p65 fused to eGFP in order to investigate the localization of the normal and the mutants RELA/p65. In both cell lines, wild-type RELA/p65 was mostly present in the cytoplasm. In contrast, the RELA/p65^H86N^ appeared to be less abundant but present in both compartments, whereas the RELA/p65^R329X^ completely miss localized, with a sequestration in the nucleus (Fig. 2B).

Immunoblotting of nuclear and cytoplasmic extracts from activated T-cell blasts indicated cytoplasmic localization of RELA/p65 wild-type and/or RELA/p65^H86N^ protein (65 kDa band) in healthy donors and P1 (Fig. S6A). In PMA-ionomycin stimulated cells, the RELA/p65 protein was detectable in nuclear cell extracts from all tested controls and patients. In extracts from P3, cytoplasmic and nuclear localization of RELA/p65^R329X^ was observed both in stimulated and non-stimulated conditions (Fig. S6A).

We then performed electrophoresis mobility supershift assays to test the RELA/p65 proteins binding to a κB consensus sequence. In unstimulated conditions, RELA/p65 binding was specifically observed with the nuclear extract of P3 (RELA/p65^R329X^) activated T cell blasts in contrast to the extracts from P1 (RELA/p65^H86N^) and a healthy donor (Fig. S6B). After PMA-ionomycin stimulation, binding of RELA/p65 was observed in all patients and control samples. Of note, two specific RELA/p65 complexes were observed for P3’s samples.

Together our analysis indicates an aberrant presence of RELA/p65 in the nucleus of the patients’ cells both at steady state and after stimulation. In particular the RELA/p65^R329X^ truncated protein is present in the nucleus, where it is able to bind to NF-κB consensus sequences on DNA.

### Functional characterization of the RELA/p65 mutants

In resting lymphocytes RELA/p65 is sequestered in the cytoplasm by complexes containing IκB-α. Furthermore, the *IκB-α* gene is a known RELA/p65 target indicating an auto-regulatory loop of NF-κB activity(Sun et al. 1993). Immuno-blotting using an antibody specific to IκB-α of total cell extracts from HEK293T cells expressing the different RELA/p65 proteins revealed that especially expression of the wild-type RELA/p65 protein induced an increased IκB-α expression (Fig. 1D). Since IκB-α expression was different in cells expressing RELA/p65 wild-type, RELA/p65^R329X^ and RELA/p65^H86N^ mutant proteins we investigated the protein abundance in cells co-expressing IκB-α and RELA/p65 mutant or wild-type proteins. We observed that co-expression of IκB-α with the different RELA/p65 proteins increased the abundance of all ectopically expressed RELA/p65 proteins, especially of the RELA/p65^H86N^ mutant protein. Interactions of the mutant RELA/p65 proteins with IκB-α and NF-κB1/p50 were confirmed by co-immunoprecipitation (Fig. 1B). Moreover, ectopically expressed p65 mutants are able to form heterodimers with p65 ^WT^ and form homodimers (Fig. S7).

To investigate the stability of the RELA/p65 mutant protein we ectopically expressed eGFP-tagged RELA/p65 mutants and wild-type protein in HEK293T cells(Fig.S89A). We analyzed RELA/p65-mediated transcriptional activation of ectopically expressed RELA/p65 mutant protein using an Ig-κ-luciferase reporter assay. The RELA/p65^H86N^ and the RELA/p65^R329X^ mutants were unable to induce a significant transcriptional activity of the Ig-κ-luciferase reporter gene (Fig. 3A). To determine whether the RELA/p65 mutant proteins interfere negatively with the transcriptional activation of RELA/p65 wild-type protein we ectopically co-expressed them in a 50/50 ratio. No significant negative interference of both mutant proteins was observed. To investigate whether the RELA/p65^R329X^ protein can form heterodimers with RELA/p65 wild-type protein we performed co-immune-precipitation experiments of ectopically expressed HA-tagged and eGFP-RELA/p65 fusion proteins. All RELA/p65 mutants were able to form the homo-dimer as well as the RELA/p65 wild-type and RELA/p65 mutant heterodimer (Fig. S7).

**Figure 3.**
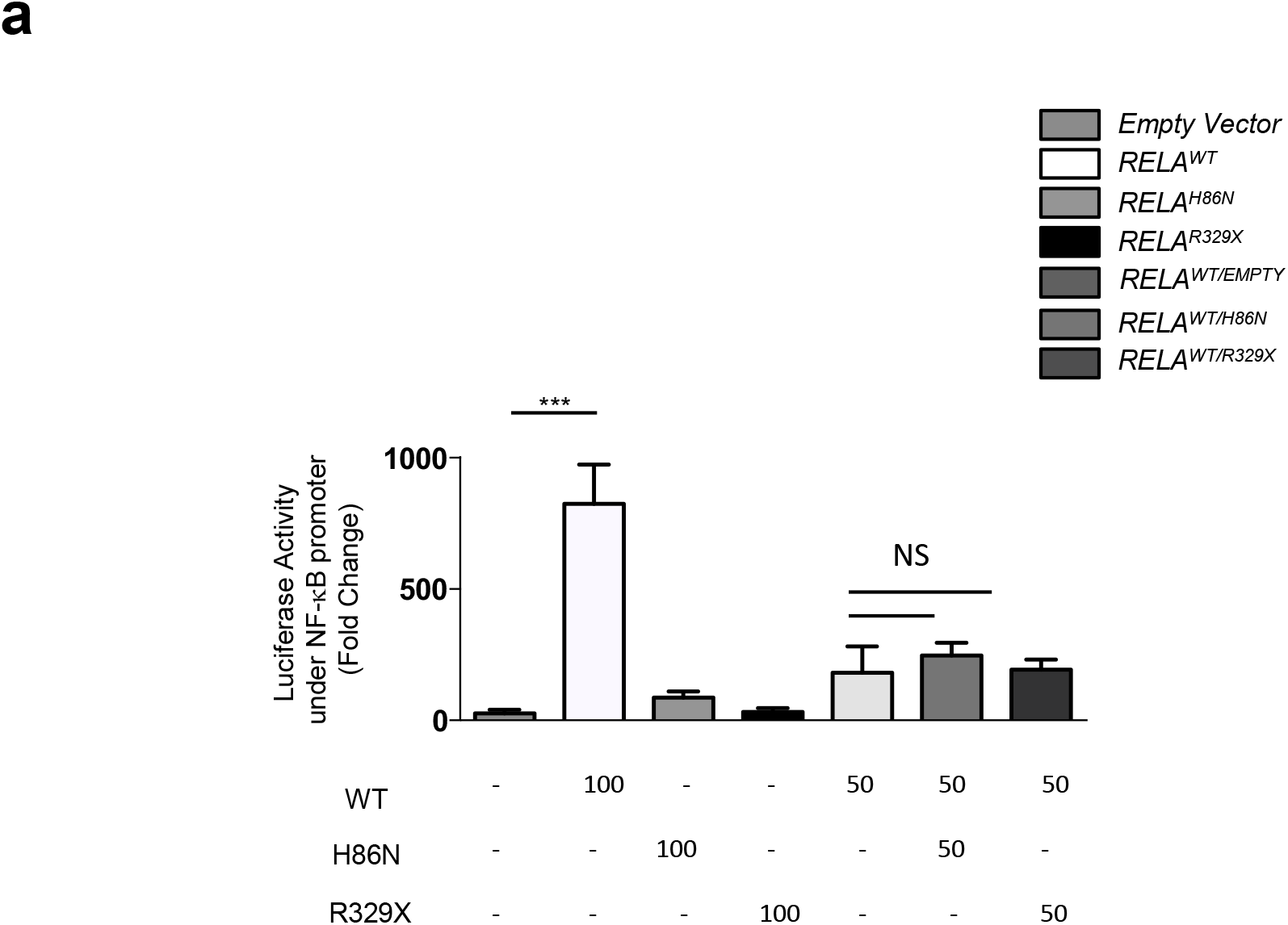
Ectopic expression and transcriptional activity of RELA/p65 proteins. Luciferase acitivity assay in HEK293T cell line stabely expressing a NF-κB consencus sequence reporter construct. with indicate ratio of co-transfected expression vectors. Depicted is the mean of 4 independent experiments with SD. Unpaired student’s t-test, *P*<0.05.

Together, these results suggest that the RELA/p65 mutant proteins are transcriptionally inactive on an Ig-κ-promoter but participate in different protein complexes.

### SLE patients’ lymphocytes display positive ISG signature

Expression of 5 ISGs was assessed on patients’ RNA obtained from PBMCs and *in vitro* activated T-cell blasts as previously described (Jeremiah et al. 2014). Expression of all 5 ISGs was significantly increased in PBMCs from the patients P1 and P3 as compared to healthy controls (Fig. 4A). Since infections may also result in positive ISGs signature, we repeated the experiments on *in vitro* activated T-cell blasts. We confirmed a positive ISGs signature in both patients (Fig. 4b). Moreover, ISG expression in PBMCs and activated T-cell blasts from P1 was comparable to that of a STING patient (activating V155M *TMEM173* mutation), used as a positive control (Fig. 4A and B) and the plasma IFN-α concentration was dramatically elevated (3059 fg/mL) (Fig. 4C). This demonstrated a strong type-I IFN dysregulation, in line with her active and severe phenotype. A milder elevation of the IFN-α plasma concentration (72 fg/mL) was observed in P3 while being in clinical remission (Fig. 5C).

**Figure 4.**
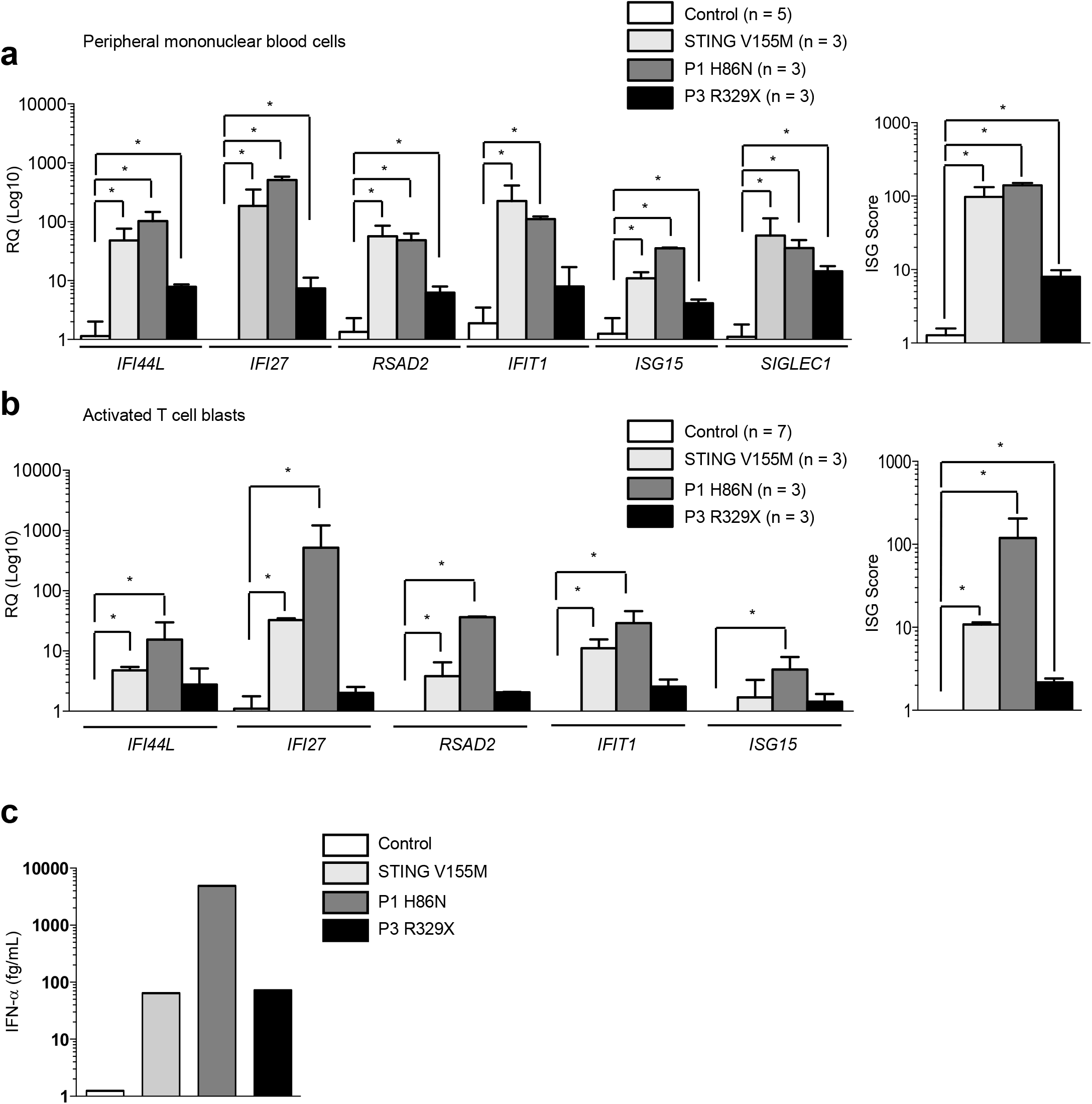
Increased expression of ISGs and IFN@ plasma concentration SLE patients. Representative quantitative RT-PCR analysis of ISGs (**a**) in Peripheral Blood Mononuclear Cells (PBMC), (**b**) activated T-cell blasts derived from PBMCs. (**c**) IFN-α concentration measured by digital ELISA (Simoa) analysis. Unpaired student’s t-test, \**P*<0.05; mean±SD.

**Figure 5.**
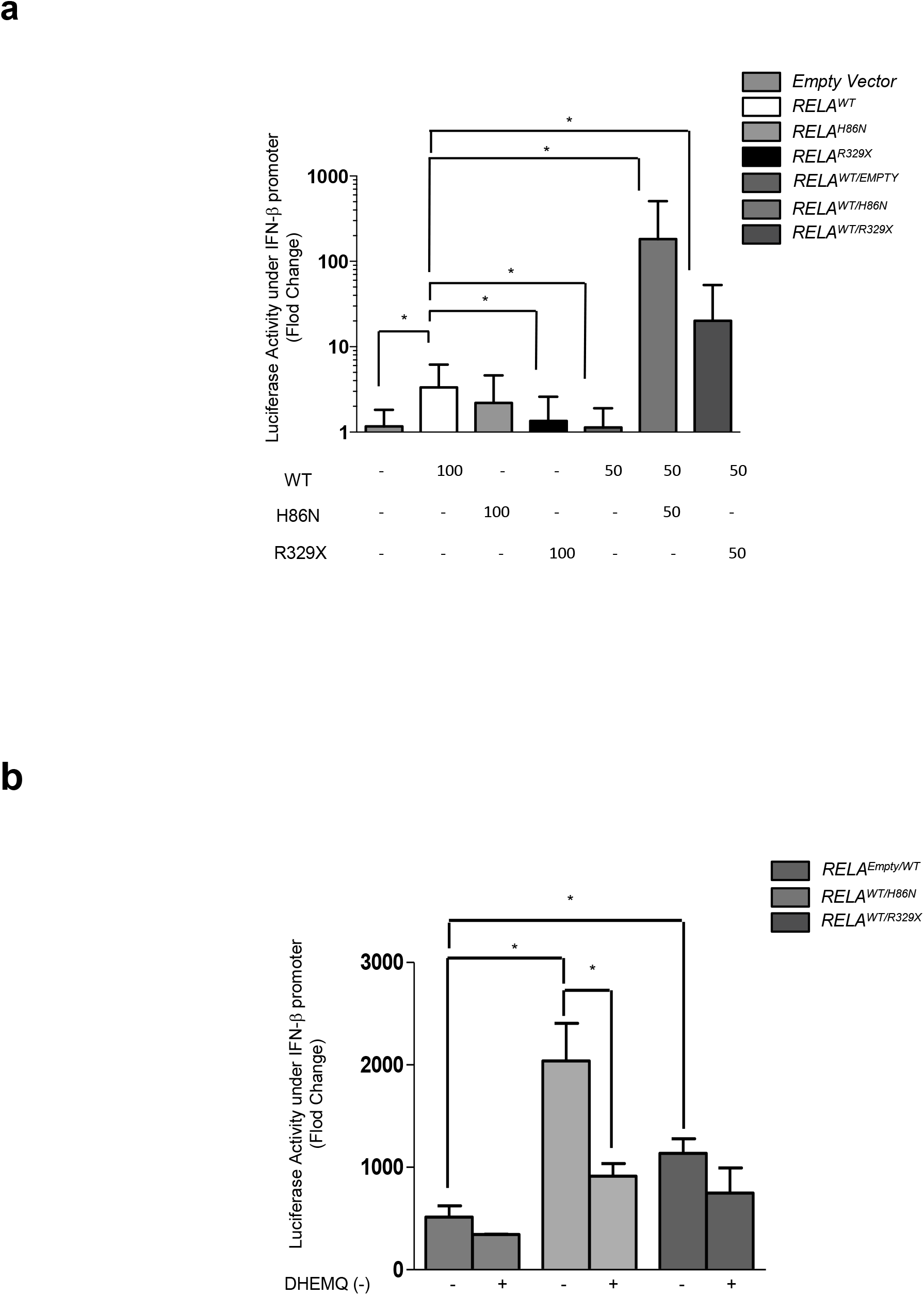
Ectopic expression and transcriptional activity of RELA/p65 proteins on interferon-β promoter and before and after DHMEQ (-) treatment. (**a**) Luciferase assay in HEK293T cell line under the control of interferon-β promoter with indicate ratio of co-transfected plasmids. Depicted is the mean of 3 independent experiments with SD. (**b**) Luciferase assay in HEK293T cell line under the control of interferon-β promoter with indicate ratio of co-transfected plasmids. Treatment with DHMEQ (-) has been done for 1 hour at a concentration of 1 μM. Depicted is the mean of 3 independent experiments with SD.

### RELA mutations leads to an interferon type I activity

In order to investigate the role of the RELA mutants in interferon pathway, we ectopically expressed eGFP-tagged RELA/p65 mutants and wild-type protein in HEK293T cells carrying a Luciferase Firefly reporter under the control of the IFN-β promoter cells expressing eGFP positive cells were sorted and luciferase activity was analyzed (Fig. S8B). The RELA/p65^H86N^ mutant, but not the RELA/p65^R329X^ mutant, was able to induce a significant transcriptional activity of the luciferase reporter gene (Fig. S8B). To determine whether the RELA/p65 mutant proteins interfere negatively or positively with the transcriptional activation of RELA/p65 wild-type protein in IFN type I regulation, we ectopically co-expressed them in a 50/50 ratio. The transfection of the empty vector with wild-type vector is accompanied with a reduced luciferase activity, thereby demonstrating that the amount of WT vector was not saturating (Fig. 5A). On the contrary, co-expression of wild-type RELA/p65 and RELA^H86N^/p65 or RELA^R329X^/p65 significantly increased the luciferase activity. This is indicating that co-expression of the wild-type and mutant RELA/p65 proteins increases transcriptional activity on IFN-β promoter (Fig. 5A).

We thus ectopically expressed eGFP-tagged RELA/p65 mutant and wild-type proteins in HEK293T cell and performed ISGs signatures after cell sorting of the transfected cells. We observed a mild but significant ISG score in cells expressing the RELA/p65^H86N^ and RELA/p65^R329X^ mutants (ISG score of 2.3 and 3.8 respectively) (Fig. S8C).

### DHMEQ(-): a possible treatment for RELA patients

Dehydroxymethylepoxyquinomicin (DHMEQ) is a newly developed compound that inhibits NFκB activation and is reported to ameliorate animal models of various inflammatory diseases without significant adverse effects (Shimizu et al. 2012). It binds the RHD domain and it works at nuclear translocation of NF-κB (Ariga et al. 2002).

We ectopically expressed eGFP-tagged RELA/p65 mutants and wild-type protein in HEK293T cells carrying a Luciferase Firefly reporter under the control of the IFN-β promoter. After cell sorting, following the protocol presented (Fig. S9), we treated these cells with DHEMQ (-) for 1 hour at 1 μM. As we observed previously, the combination of both wild-type and mutant RELA/p65 proteins resulted in an important increase of of the luciferase activity (Fig. 5B). The treatment with DHMEQ strongly decreased transcriptional activity on IFN-β promoter of co-expressed wild-type and mutant RELA/p65 proteins whereas transcriptional activity of wild-type RELA/p65 protein barely affected (Fig. 5B). B cells of the patient *RELA*^R329X^ have been immortalized by Epstein Barr Virus (B-EBV) and have been treated with DHMEQ (-) for 1 hour at 1 uM. A decrease of the ISG score by half has been observed (Fig. S10).

## Discussion

Monogenic causes of SLE are increasingly reported, with greater prevalence among male patients and those with severe early-onset disease and syndromic lupus. These rare monogenic SLE cases can help to understand SLE pathogenesis and identify therapeutic targets. To date, clearance of defective apoptotic bodies (complement deficiencies(Macedo and Isaac 2016)), B cell-defective apoptosis (Protein Kinase Cδ Deficiency)(Belot et al. 2013), early B cell differenciation defect (*IKZF1* mutations)(Hoshino et al. 2017) and disturbance of nucleic acids metabolism (Type I interferonopathies, Dnase1L3 mutation)(Lee-Kirsch 2017; Al-Mayouf, Abanomi, and Eldali 2011) are the main known causes associated with a susceptibility to SLE. Herein, we identified mono-allelic *RELA* mutations leading to dysregulation of the NF-κB pathway in three patients with severe early-onset and/or familial SLE and upregulation of the type-I interferon pathway. These mutations resulted in the expression of mutant RELA/p65 proteins, a RELA/p65^H86N^ in P1 and P2 and a truncated RELA/p65^R329X^ in P3. RELA/p65 is a transcription factor of the NF-κB family. When bound to IκB-α, RELA/p65 is sequestered in the cytosol. Upon phosphorylation by the IKK complex, IκB-α is degraded by the proteasome, allowing nuclear translocation of RELA/p65, where it mediates its transcriptional activity.

Recently, *RELA*/RELA/p65 haploinsufficency has been described in patients presenting with either muco-cutaneous ulcers(Badran et al. 2017) or lymphoproliferation with autoimmune cytopenia(Comrie et al. 2018). In contrast the *RELA* mutation identified in the SLE patients results in expression of mutant (RELA/p65^H86N^ or RELA/p65^R329X^, respectively) proteins. Based on the inheritance and expression of the mutant RELA/p65 proteins in the SLE patients, we considered that they behave as dominant negative or gain of function mutants interfering with NF-κB regulation and/or function. Accordingly, by confocal microscopy analyses, we observed a spontaneous RELA/p65 nuclear localization in unstimulated patients’ cells. An increased nuclear localization was also observed after stimulation of the NF-κB pathway, confirming a global RELA/p65 dysregulation. These analyses were further confirmed by western blot experiments evidencing the presence of the RELA/p65^R329X^ truncated mutant (consisting mostly of the RHD) in nuclear extracts of both stimulated and unstimulated P3’s cells. In addition, the RELA/p65^R329X^ mutant was able to bind to a NF-κB DNA consensus sequence as well as to other protein partners e.g. NF-κB1/p50, RELA/p65^WT^ and RELA/p65^R329X^ itself. Molecularly, we could not distinguish the RELA/p65^H86N^ in P2’s cells from the RELA/p65^WT^ protein. Analysis of activated T-cell blasts from P1 and P2 suggests normal RELA/p65 protein expression as compared to control activated T-cell blasts. Nevertheless, ectopically expressed RELA/p65^H86N^ protein was less stable than RELA/p65^WT^, indicating that the amino acid substitution might impact the RELA/p65^H86N^ protein integrity. Co-immunoprecipitation analysis demonstrated interaction of RELA/p65^H86N^ with @IkB-α@ and NF-κB1/p50. Increased levels of RELA/p65^H86N^ suggests its stabilization by@IkB-α®in cells co-expressing RELA/p65 and®IkB-α. Altogether, these results suggest that the pathogenic mechanism of RELA/p65^H86N^ and RELA/p65^R329X^ involves disturbed homeostasis of NF-κB dimers.

Both mutant proteins were transcriptionally inactive in an Ik luciferase reporter assay when ectopically expressed. This was expected from the RELA/p65^R329X^ mutant lacking the transactivation domain. Interpretation of the results obtained with the RELA/p65^H86N^ mutant is uneasy considering its possible impaired integrity *in vitro*. However, the ectopic co-expression of RELA/p65^WT^ with RELA/p65 mutant proteins argues against an autosomal dominant-negative effect on transcriptional activity. Nevertheless, the consequences in the patients’ cells were similar in terms of spontaneous RELA/p65 activation and nuclear localization, suggesting a lack of control of the NF-κB pathway activation and probably a subsequent lower cellular activation threshold. Such dysregulation of the NF-κB pathway is reminiscent with the recently described loss-of-function mutations of the *TNFAIP3* gene leading to A20 haploinsufficiency (HA20)(Duncan et al. 2018; Comrie et al. 2018; Takagi et al. 2017; Zhou et al. 2016). HA20 results in a dominantly inherited, early-onset systemic inflammation and a broad spectrum of autoinflammatory and autoimmune features, especially Behcet-like disease. In a cohort of 16 patients with HA20, half of them developed auto-immune hepatitis, Hashimoto’s thyroiditis and low-titer of autoantibodies (antinuclear, anti-dsDNA, and lupus anti-coagulant), and SLE-like disease was seen in one patient(Aeschlimann et al. 2018). Interestingly, all 3 SLE patients with *RELA* mutations had oral ulcers which is a hallmark in HA20 patients. HA20 resulted in increased activation of the NF-κB pathway upon TNF-α stimulation in all instances, and in particular to an excess of RELA/p65 phosphorylation(Duncan et al. 2018) and nuclear localization(Zhou et al. 2016). Accordingly, an increased expression of RELA/p65-regulated genes, such as *IL6, TNF-α, IL1-β, IL17* or *IFN-I* was observed in most HA20 patients. Although the precise underlying molecular mechanisms remain to be established, notably to explain the variable clinical outcomes of a common monogenic deficiency, these observations pointed to a global dysregulation of the NF-κB pathway. In the present study, we did not observe elevated expression of *TNF-α* although a slightly elevated *IL6* and *IL1-β* expression at the transcriptional level in P1 and P3 respectively was observed. However, this was not confirmed by the production of the corresponding cytokine in the patients’ plasma (data not shown). The consequences of the present *RELA* mutations leading to the expression of mutant RELA/p65 proteins result in a different spectrum of dysregulation as compared with *RELA* or *TNFAIP3* haploinsufficiencies.

As reported in most SLE patients, we found a marked increase of ISG expression in the SLE patients with *RELA* mutations, likely resulting from the type-I IFN overproduction detected in the patients’ sera. Whether IFN-I overproduction is the cause or the consequence of SLE remained a matter of discussion so far. But one could not exclude a direct effect of the RELA/p65 activation in the patients’ cells. Indeed, a RELA/p65-IRF5 interaction has been described to target inflammatory genes in macrophages(Saliba et al. 2014). We are now showing that co-expression of the mutant and WT RELA/p65 in HEK293T cell line leads to the expression of a reporter gene under the control of the INF-β promoter. This experiment indicates that the mutant RELA/p65 proteins hijacke and protmote the RELA/p65 transcriptionnal activity towards the type I IFN genes whereas no interference was observed on Ig-κ consensus sequences. Of note, P1 exhibited a persistent hypogammaglobulinemia long after anti-CD20 therapy, although her immunoglobulin levels were in the normal range before the treatment. This could possibly be the consequence of a reduced activity of the RELA/p65 on the Immunoglobulin genes promoters. A similar adverse effet has been reported in a patient with a spondylenchondrodysplasia due to biallelic ACP5 mutations(Briggs et al. 2016). Although these obsersations could rely on different mechanisms, they suggest that anti-CD20 therapy should be used with caution for treating autoimmune manifestations in a subset of patients presenting with monogenic SLE or interferonopathies.

In conclusion, heterozygous *RELA* mutations are a new monogenic cause of early-onset and/or dominantly inherited familial SLE. They led to the expression of mutant RELA/p65 proteins and a disturbed homeostasis of NF-κB dimers associated with an overall dysregulated lymphocyte activation. This study is thus providing evidence for the involvement of the NF-κB pathway in the pathogenesis through an overexpression of the type I IFN genes, and it is opening future avenues to identify new therapeutic targets. Currently, no NF-κB inhibitors have progressed to the clinic for the treatment of autoimmunity, but a variety of promising approaches targeting multiple stages of the NF-κB pathway are currently being explored. Some of them are targeting the IKK complex, exerting their activity via interaction with the conserved adenosine triphospate (ATP) binding site of IKK molecules (Palanki et al. 2003). Other strategies regard the targeting of IκB-α. In particular, the use of IκB-α super-repressor (IκBαSR), a mutated form of IκB-α sequestration in the cytoplasm of the cells and inhibition of its activity(Taylor et al. 1999). Targeting the DNA binding domain with some drugs, such as DHMEQ(-), which binds to specific cysteine residues of Rel family, could represent another strategy (Taylor et al. 1999). It has been shown that DHMEQ reduces the serum levels of anti-dsDNA, anti-nucleosome and anti-histone antibodies in mice with pristane-induced lupus(Qu, Bian, and Xu 2014). More generally, DHMEQ is reported to ameliorate animal models with various inflammatory diseases without significant adverses effect(Shimizu et al. 2012). Our study suggests that DHMEQ (-) could represente a promising therapeutic strategy for arly-onset SLE patients associated to disturbed RELA/p65 protein function caused by *RELA* mutations.

### Patients and Methods

We initated a project (Lumugene) aiming to identify monogenic causes predisposing to SLE in sporadic early-onset or familial SLE. We gathered a cohort of more than 90 patients with pediatric onset and/or familial SLE. Peripheral blood and tissue samples were obtained from the patients or their relatives after the provision of informed written consent in accordance to the Declaration of Helsinki. This study received the agreement of the Ile de France ethic committee (CPP IDF2 DC-2014-22722015-03-03 AF) and the samples collection has been declared to the French ministry of research with the reference DC 2014-2272

### Statistical analysis

Analyses were performed with PRISM software (version 7 for Macintosh, GraphPad Inc.) or R. Statistical hypotheses were tested if not otherwise indicated using unpaired student’s t-test. Statistical analysis of RELA/p65 nuclear localization analyzed by immunofluorescence was performed with a Mann-Whitney Wilcoxon test. Statistical analysis for ectopically expressed HA-RELA/p65 and IκB-α data normalized on α-tubulin expression was performed with a paired student’s t-test on log2 transformed data. A P value less than 0.05 was considered significant for all statistical tests; *, P < 0.05.

Further details are provided in the Supplemental Methods

## Acknowledgements

The authors would like to thank Dr E. Bal for providing the plasmids for the NF-κB luciferase reporter assay. The pEBB HA RelA (Addgene plasmid # 74892) and GFP-RelA (Addgene plasmid # 23255) plasmids were gifts from Ezra Burstein & Colin Duckett and Warner Greene, respectively. We would like to thank E. Dandelot, Dr A. Guérin and the cell imaging platform of the IMAGINE Institute (M. Garfa-Traoré and Nicolas Goudin) for technical advice. D.D. Thanks ImmunoQure AG for sharing of antibodies used in the Simoa assay. Thanks Nicolas Cagnard from Imagine Institute’s Bioinformatics Facility for advice on statistical analysis. We acknowledge the use of the bioresources of the Necker Imagine DNA biobank (BB-033-00065).

S.K. is a Centre National de la Recherche Scientifique staff researcher. The study was supported by the Institut National de la Santé et de la Recherche Médicale (INSERM) and by a government grant managed by the Agence National de la Recherche as part of the “Investment for the Future” program (ANR-10-IAHU-01), the Ligue Contre le Cancer – Comité de Paris, Fondation ARC pour la recherche sur le CANCER, the Centre de Référence Déficits Immunitaires Héréditaires (CEREDIH) and the Agence National de la Recherche (ANR-14-CE14-0026-01 “Lumugene” to FRL and ANR-15-CE15-0020 ANR-PIKimmun to S.K.). L.B. was a recipient of an Imagine institute PhD international program supported by the Fondation Bettencourt Schueller. L.B. was also supported by the EUR G.E.N.E. (reference #ANR-17-EURE-0013) and is part of the Université de Paris IdEx #ANR-18-IDEX-0001 funded by the French Government through its “Investments for the Future” program. H.L. received fellowships from the French Ministry of research at the University Paris Descartes and from IMAGINE Institute. NGS was supported by a funding program of the Fondation maladies rares (AAP2012 High throughput sequencing). D.D. acknlowedeges the ANR (grant CE17001002) for funding support. The authors declare no competing financial interests.

## Author contributions

F.RL. and S.K. conceived study, planned experiments, analyzed data and wrote the manuscript and L.B., H.L. and M.C. participated in writing of the manuscript. L.B. and N.J. identified *RELA* mutations. L.B., H.L., M.C. planned, performed experiments and analyzed data with help from S.J., MC.S., A.MC., M.M., C.U., L.C., A.B., M.C. D.D. and V.B. performed digital elisa.

C.P. produced immune phenotyping.

M.P. performed WES, and M.B. performed bioinformatics analysis of exome data.

B.BM., R.S., C.B., M.H. provided patient care and clinical data.

## Supplementary Material and methods

### Cell culture

Peripheral Blood Mononuclear Cells (PBMC) were isolated by Ficoll-Paque density gradient centrifugation and washed twice with PBS. Activated T-cell blasts were obtained by stimulating 1×10^6^ cells per mL with staphylococcal enterotoxin (SEE) 0.1 μg/mL (Toxin Technology) in Panserin 401 (PAN Biotech) supplemented with 10% human AB serum (Biowest), 1x penicillin - streptomycin (P/S) (Invitrogen), 1% glutamine. After 3 days of activation, viable cells were separated by Ficoll centrifugation washed twice with Panserin and then cultured in Panserin supplemented with 10% human AB serum, 1% P/S and 100 U/ml interleukin 2 (IL-2). IL2 was refreshed every 2 to 3 days.

### DNA and RNA preparation

DNA was isolated from the red blood cell pellet, obtained after Ficoll preparation of fresh blood samples, using the QiAamp DNA blood Mini kit. PCR was performed using Go Taq Flexi DNA polymerase (Promega) following manufacturer’s instructions and with the following pairs of olinucleotides: forward_Mut A 5’-ggcgagaggagcacagatac −3’, reverse_Mut A 5’-cctgggtccagaaaggagta −3’ and forward_Mut B 5’-ggactgggaaagccagagag - 3’, reverse_Mut B 5’-cccaggagtcttcatctcca −3’. PCR products were purified on sephadex G-50 columns (GE healthcare). Purified PCR products were used as template for Sanger sequencing reaction using BigDye Terminator v3.3 cycle sequencing kit (Applied Biosystems) and 3500XL genetic analyser (Applied Biosystems). Forward primers were used for sequencing reaction. All sequences data were analyzed using ApE and 4peaks software.

Total RNA was isolated from PBMCs, activated T-cell blasts line using the RNeasy mini Kit following manufacturer’s instructions (QIAGEN RNeasy mini kit) and cDNA was prepared using the Quantitect Reverse Transcription Kit according to manufacturer’s instructions including DNAse treatment to deplete genomic DNA, according to the manufactures instructions (RNase-Free Dnase Set from QIAGEN). PCR was also performed on cDNA using Go Taq Flexi DNA polymerase (Promega) with the following pair of olinucleotides: forward_Mut A 5’-gcgagaggagcacagatacc-3’, reverse_Mut A 5’-tggtcccgtgaaatacacct −3’ and forward_Mut B 5’-caagtggccattgtgttccg-3’, reverse_Mut B 5’-ccccttaggagctgatctga-3’. Forward primers were used for sequencing reaction.

### Gene expression analysis

The expression of a set of 6 interferon-stimulated genes (ISGs) in PBMCs and activated T-cell blasts was assessed by quantitative RT-PCR using TaqMan Gene Expression Assays. Expression of ISGs was normalized to GAPDH. The following TaqMan Gene Expression Assays obtained from ThemoFisher Scientific were used: *IFI27* - Hs01086370_m1, *IFI44L-* Hs00199115_m1, *IFIT1* - Hs01675197_m1, *RSAD2* - Hs00369813_m1, *SIGLEC1-* Hs00988063_m1, *ISG15* – Hs01921425_s1, *GAPDH-* Hs03929097. Real Time quantitative PCR was performed in duplicate using the LightCycler VIIA7 System (Roche). Relative quantification (RQ) represents the fold change compared to the calibrator (*GAPDH*). RQ equals 2^-ΔΔCt^ where ΔΔCt is calculated by (*Ctt_ar_g_e_t* _-_ CT*GAPDH*) test sample - (Ct*target* - Ct*GAPDH*) calibrator sample. Each value is derived from three technical replicates.

### Confocal Microscopy

Activated T-cell blasts (3×10^6^) were stimulated in 3 ml of complete Panserin with either PMA (20 ng/mL; Sigma-Aldrich) and ionomycin (1 μmol/L) or 10ng/ml TNF-α (Invitrogen) for 15 minutes and 30 minutes at 37°C. Cells were coated on a poly-L-lysine matrix then fixed 20min with PFA 4% at room temperature (RT). Cells were permeabilized with PBS + 0.1% Triton X-100 10min at RT. To block unspecific staining samples were blocked for 1h with PBS + 5% BSA at RT before staining. Antibodies were diluted in PBS + 5% BSA. The primary antibodies used included rabbit monoclonal anti human NF-κB/p65 (E379) C terminal (1/200; Abcam) and a mouse monoclonal anti human NF-κB/p65 (F-6) N terminal (1/100; SantaCruz). Samples were incubated overnight at 4°C. For immunofluorescence, the following secondary antibodies were used: goat anti-rabbit IgG labelled with Alexa 488 and donkey anti-mouse labelled with Alexa 555 (1:1000; Invitrogen) 1h at RT in the dark. Slides were counterstained after 3 washes of PBS with 0.3 μg/mL 4,6-diamidino-2-phenylindole (Sigma-Aldrich).

Images including Z-stacks were acquired on a Leica SP8 gSTED with an objective with x63 magnification. Nuclear localization of p65 was quantified by ImageJ software.

HEK293T cells and HacaT cells have been transfected using Lipofectamine 2000 (Thermo Fischer Scientific) following manufacturer’s instructions. Images have been acquired on a Leica SP8 gSTED, evaluating the EGFP fluorescence.

### Immunoblotting of patient cells

Protein samples were extracted in RIPA buffer (Bio-Rad-ThermoFisher) supplemented with Complete Protease Inhibitor Cocktail (Roche). Equal amounts of extracted protein (10 μg) were loaded and run alongside Chameleon Duo Pre-stained Protein Ladder (LI-COR) on a NuPAGE 4–12% Bis-Tris Gel (Invitrogen) and transferred onto nitrocellulose membranes using iBlot2 (Invitrogen). The membranes were incubated with anti-βactin (13E5 1:10,000 rabbit polyclonal; Cell Signaling) or anti-p65 (F-6 1:1000; mouse monoclonal Santa Cruz and 1: 2,000; E379 rabbit monoclonal; Abcam) primary antibodies. Immuno-reactive bands detected following incubation with IRDye 680 RD anti-rabbit antibody (1:10000; LI-COR) and IRDye 680 RD anti-mouse antibody (1:10000; LI-COR) by using the CLx Odyssey Infrared Imaging System (LI-COR). Quantification of immunoblots was performed by using the Image Studio Lite software version 5.0 (LI-COR).

### Expression plasmids

Site directed mutagenesis using GeneART™ Site-Directed Mutagenesis System (Thermo Fischer Scientific) was performed following manufacture’s instruction on the *RELA*^WT^ expression constructs pEBB HA RelA (addgene #74892) and GFP-p65 (addgene #23255). The following oligonucleotide pairs were used for mutagenesis: forward *RELA*^H86N^ 5’-CCTCCTCACCGGCCTAACCCCCACGAGCTTG-3’, reverse *RELA*^H86N^ 5’-CAAGCTCGTGGGGGTTAGGCCGGTGAGGAGG-3’, forward *RELA*^R329X^ 5’-CCCCGGCCTCCACCTTGACGCATTGCTGTGC-3’, reverse *RELA*^R329X^ 5’-GCACAGCAATGCGTCAAGGTGGAGGCCGGGG-3’. The *RELA* sequence of all the plasmids were confirmed by DNA Sanger sequencing. *IκB-α* expression plasmid was a courtesy of Robert Weil (Pasteur Institute, Paris).

IgK-luc (Firefly luciferase) and Renilla-Luciferase constructs were a courtesy of Elodie Bal (IMAGINE institute, Paris).

### Ectopic expression of RELA mutant protein

All transfection of HA-65 mutants or wild type expressing plasmids into HEK293T cells were carried out using lipofectamine 2000 (Invitrogen) according to the manufacturer’s instructions. Cells were lysed 48h after transfection for subsequent assays.

### NF-κB luciferase reporter assay

NF-κB reporter Luc-HEK293T recombinant cell line (BPS Bioscience) were transfected using Lipofectamine 2000(Thermo Fischer Scientific), following manufacturer’s instructions. After 24h, EGFP cells were sorted and seeded in 96-well plates. Luciferase assay was performed using the Luciferase assay system (Promega) following manufacturer’s instructions. Luminescence was analyzed with a VictorX4 plate-reader (PerkinElmer).

### IFNbeta luciferase reporter assay

HEK293T have been transfected with a plasmid containing the luciferase under the control of the IFN-β promoter. The plasmid was a gift from Dr. Nicolas Manel. HEK293T have been also transfected with RELA WT, RELA H86N, RELA R329X and a ratio of 50/50 RELA H86N/ RELA WT and RELA R329X/RELA WT. Transfection experiment has been carried out using lipofectamine 2000 (Thermo Fischer Scientific) following manufacturer’s instructions. eGFP positive cells have been cell sorted 24H after transfection and have been seeded in in 96-well plates. Luciferase assay was performed using the Luciferase assay system (Promega) following manufacturer’s instructions. Luminescence was analyzed with a VictorX4 plate-reader (PerkinElmer).

### Immunoblotting of transfected cell lines

For immunoblot analysis cell lysates of 1×10^6^ HEK293T cells were prepared with RIPA buffer 50 mM Tris pH7.4, 1% Triton X100, 0.5% sodium deoxycholate, 0.1% SDS, 150 mM NaCl, 2 mM EDTA with HALT protease inhibitor cocktail (Pierce). To ensure equal loading, protein concentrations were assessed via BCA^™^ protein assay kit (Thermo Fischer Scientific) before protein loading on SDS/PAGE 4-12% pre-casted polyacrylamide gels (Thermo Fischer Scientific). The antibodies NF-κB p65 (C20; recognizing a C-Terminal epitope of p65, sc-372), NF-κB p65 (F6; recognizing a N-Terminal epitope of p65; sc-8008), Lamine A/C (E-1; sc**-376248)** and anti-IκB-α (H-4, sc-1643) were obtained from Santa Cruz Biotechnology. Anti-HA rabbit (**H6908**; Sigma) and anti-α-tubulin (**T5168;** Sigma) were obtained from Sigma Aldrich. All antibodies were used according to the manufacturer’s instructions. Secondary fluorescent antibodies were obtained from LI-COR and were used in immunoblotting assays. Fluorescence analysis of immunoblots were performed on digitized images using ImageStudioLite version 5.2.5 software.

### Electrophoretic Mobility Super-shift assay

Nuclear and cytoplasmic extracts for electrophoretic mobility super-shift assay were prepared using a NE-PER nuclear and cytoplasmic extraction kit (Thermo Scientific) according to manufacturer’s instructions. Cells analyzed were treated with phorbol 12-myristate 13-acetate (PMA) (20 ng/mL; Sigma-Aldrich) and ionomycin (1μmol/L; Sigma-Aldrich) for 30 min or left untreated. DNA-protein interaction mixture was performed by using nuclear extracts and the Odyssey EMSA Buffer Kit with NF-κB IRDye 700 Infrared labeled oligonucleotides 5’-AGTTGAGGGGACTTTAGGC-3’ and 3’-TCAACTCCCCTGAAAGGGTCC G-5’ (underlined nucleotides are the kB consensus binding site), following manufacturer’s instructions. Super-shift of complexes was produced by adding 1 μg of anti-p65 (F6, sc-8008) antibody to the oligonucleotide-nuclear extract mixture 20 min prior to loading. Samples were subjected to PAGE (4 % gels, under non denaturation condition) and analyzed by Odyssey CLX LI-COR. Fluorescence analysis of electrophoretic mobility super-shift assays were performed on digitized images using ImageStudioLite version 5.2.5 software.

### Immunoprecipitation

For immunoprecipitation analysis, cell lysates of 1×10^6^ HEK293T cells were prepared 100 μl of EBC lysis buffer (120 mM NaCl, 50 mM Tris [pH 8.0], 5 mM EDTA, 50 mM Hepes, and 0,5% NP-40) supplemented with protease inhibitor. After 1 h incubation in lysis buffer at 4°C, the lysates were then centrifuged for 10 min at 13,200 rpm at 4°C, and the protein concentration was determined as in the previous immunoblotting assays. Immunoprecipitation was performed with HA-Tag (C29F4, Cell Signalling) Rabbit mAb (Magnetic Bead Conjugate) according to the manufacturer’s instructions. Initial lysates and immuno-precipitated proteins were analyzed as previous immunoblotting assays.

### Simoa digital ELISA

Interferon alpha protein was measured on a single molecule array (Simoa) platform using two autoantibodies specific for interferon alpha isolated and cloned from APS1 / APECED patients (PMID: 27426947). The 8H1 antibody clone was used as the capture antibody after coating onto paramagnetic beads (0.3mg/mL), and the biotinylated (biotin/Ab ratio = 30/1) 12H5 as the detector. Recombinant IFN-α17/αI (PBL Assay Science) was used as a standard curve after cross-reactivity testing. The limit of detection was calculated by the mean value of all blank runs + 3SDs and was 0.23 fg/mL.

### Treatment with DHMEQ (-)

BEBV (B cells immortalized by Epstein Barr Virus) of the patient RELA R329X have been plated 24 hours before the treatment with DHMEQ (-) (Med Chem Express). The concentration of the drug has been established according to the manufactory instruction. A dose response curve has been determined using propidium iodide (PI). Cells have been treated for 1 hour with a concentration of 1 uM DHMEQ (-).

EGFP positive transfected cells (previosuly described) have been cell sorted 24h after transfection and have been seeded in 96-well plates. Treatment with the DHMEQ (-) has followed for 1 hour at a concentration of 1 μM of DHMEQ (-).

