## Supplementary material for "Heterozygous RELA mutations cause early-onset systemic lupus erythematosus by hijacking the NF-κB pathway towards transcriptional activation of type-I Interferon genes": Fig, S10

### Slide 1
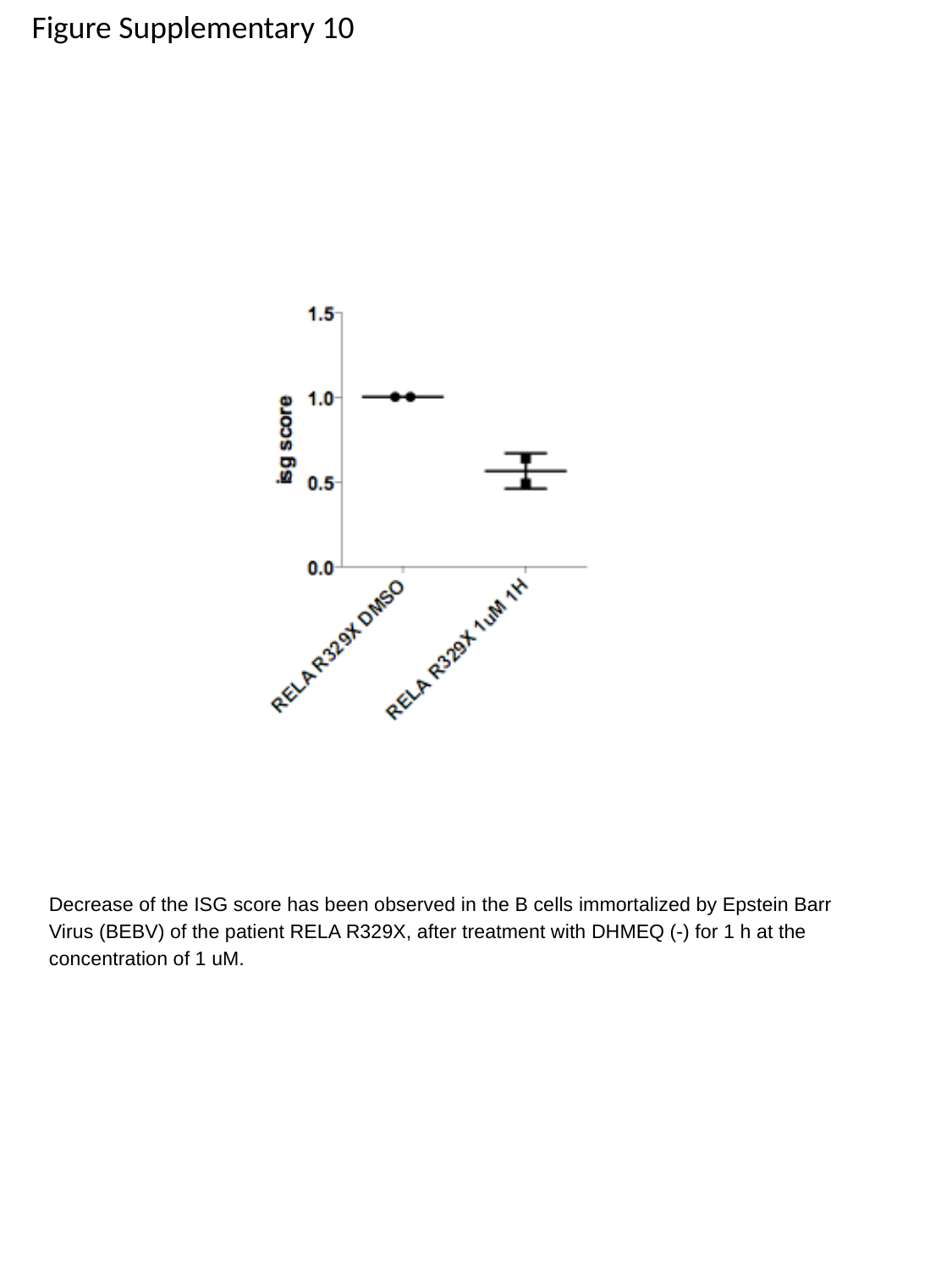

Figure Supplementary 10
Decrease of the ISG score has been observed in the B cells immortalized by Epstein Barr Virus (BEBV) of the patient RELA R329X, after treatment with DHMEQ (-) for 1 h at the concentration of 1 uM.
