## Supplementary material for "Heterozygous RELA mutations cause early-onset systemic lupus erythematosus by hijacking the NF-κB pathway towards transcriptional activation of type-I Interferon genes": Fig. S9

### Slide 1
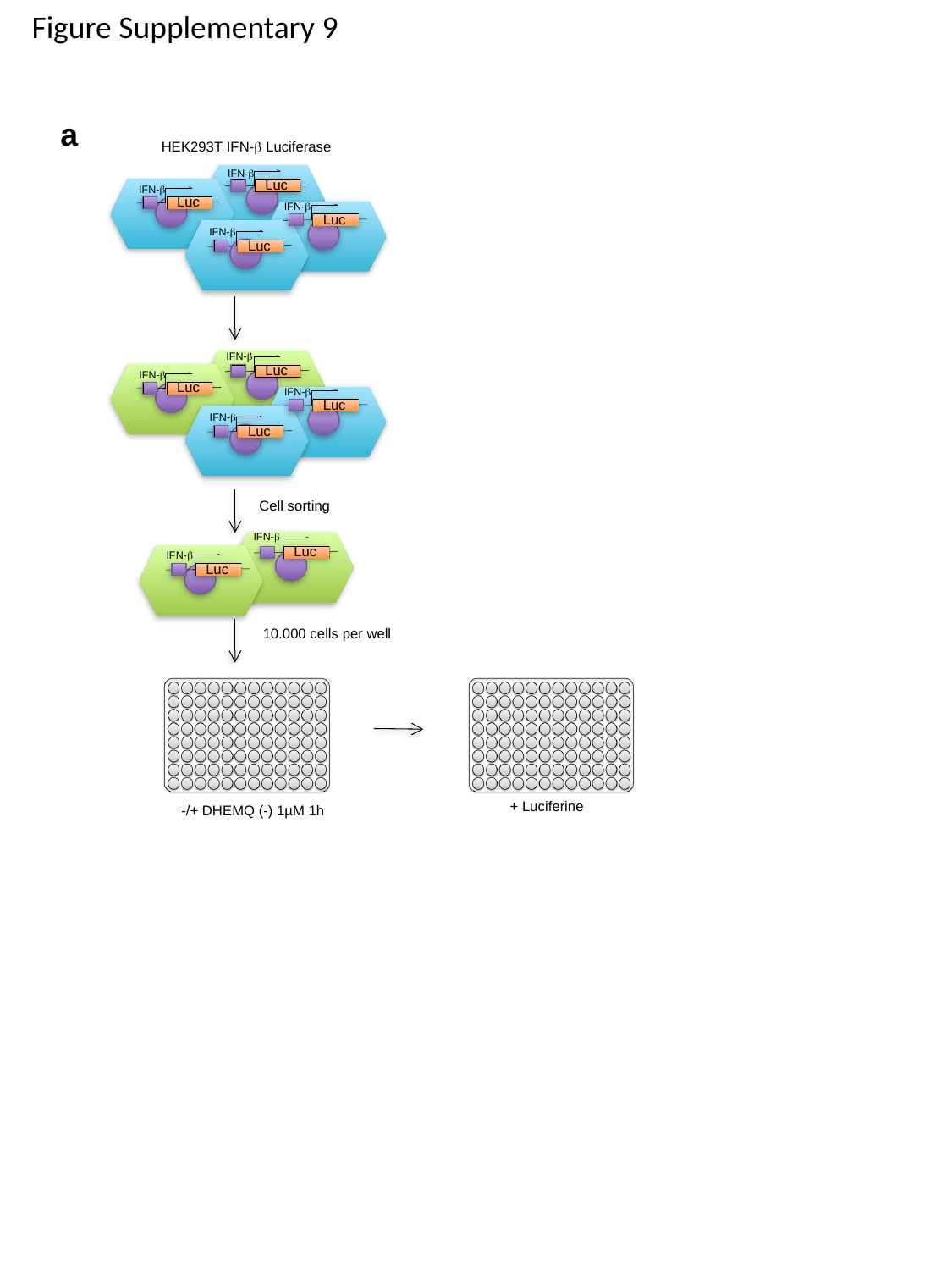

Figure Supplementary 9
a
HEK293T IFN-β Luciferase
IFN-β
Luc
Luc
IFN-β
Luc
IFN-β
Luc
IFN-β
Luc
Luc
IFN-β
Luc
IFN-β
Luc
Cell sorting
IFN-β
Luc
IFN-β
Luc
IFN-β
IFN-β
10.000 cells per well
+ Luciferine
-/+ DHEMQ (-) 1µM 1h
