## Supplementary material for "Heterozygous RELA mutations cause early-onset systemic lupus erythematosus by hijacking the NF-κB pathway towards transcriptional activation of type-I Interferon genes": Fig, S8

### Slide 1
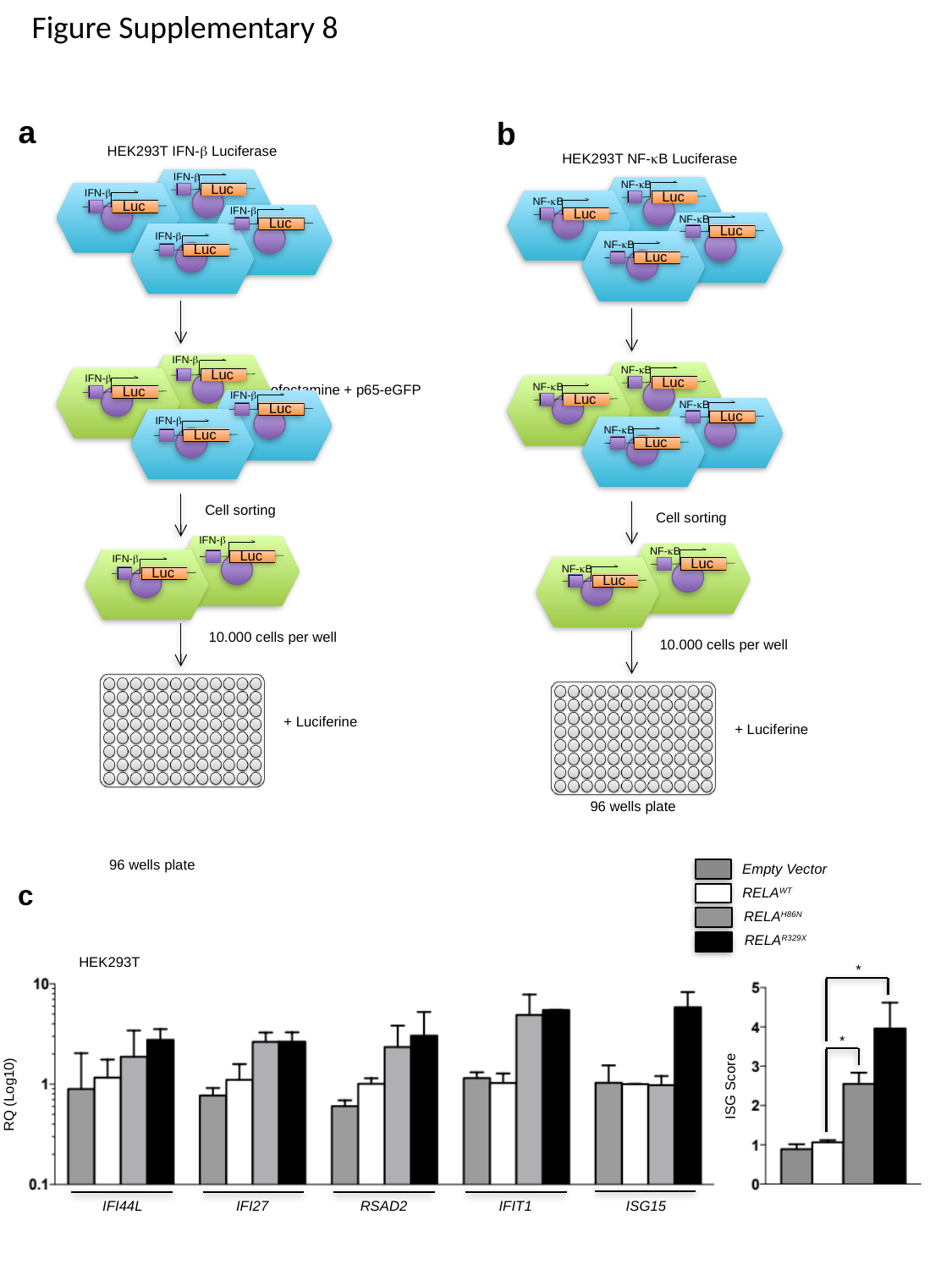

Figure Supplementary 8
a
b
HEK293T NF-κB Luciferase
NF-κB
Luc
NF-κB
Luc
NF-κB
Luc
NF-κB
Luc
NF-κB
Luc
NF-κB
Luc
NF-κB
Luc
NF-κB
Luc
Cell sorting
NF-κB
Luc
NF-κB
Luc
10.000 cells per well
+ Luciferine
96 wells plate
HEK293T IFN-β Luciferase
IFN-β
Luc
Luc
IFN-β
Luc
IFN-β
Luc
IFN-β
Luc
Luc
IFN-β
Luc
IFN-β
Luc
Cell sorting
IFN-β
Luc
IFN-β
Luc
IFN-β
IFN-β
10.000 cells per well
+ Luciferine
Lipofectamine + p65-eGFP
96 wells plate
Empty Vector
c
RELA^WT^
RELA^H86N^
RELA^R329X^
HEK293T
*
*
ISG Score
RQ (Log10)
IFI44L
IFI27
RSAD2
IFIT1
ISG15
