## Supplementary material for "Heterozygous RELA mutations cause early-onset systemic lupus erythematosus by hijacking the NF-κB pathway towards transcriptional activation of type-I Interferon genes": Fig. S7

### Slide 1
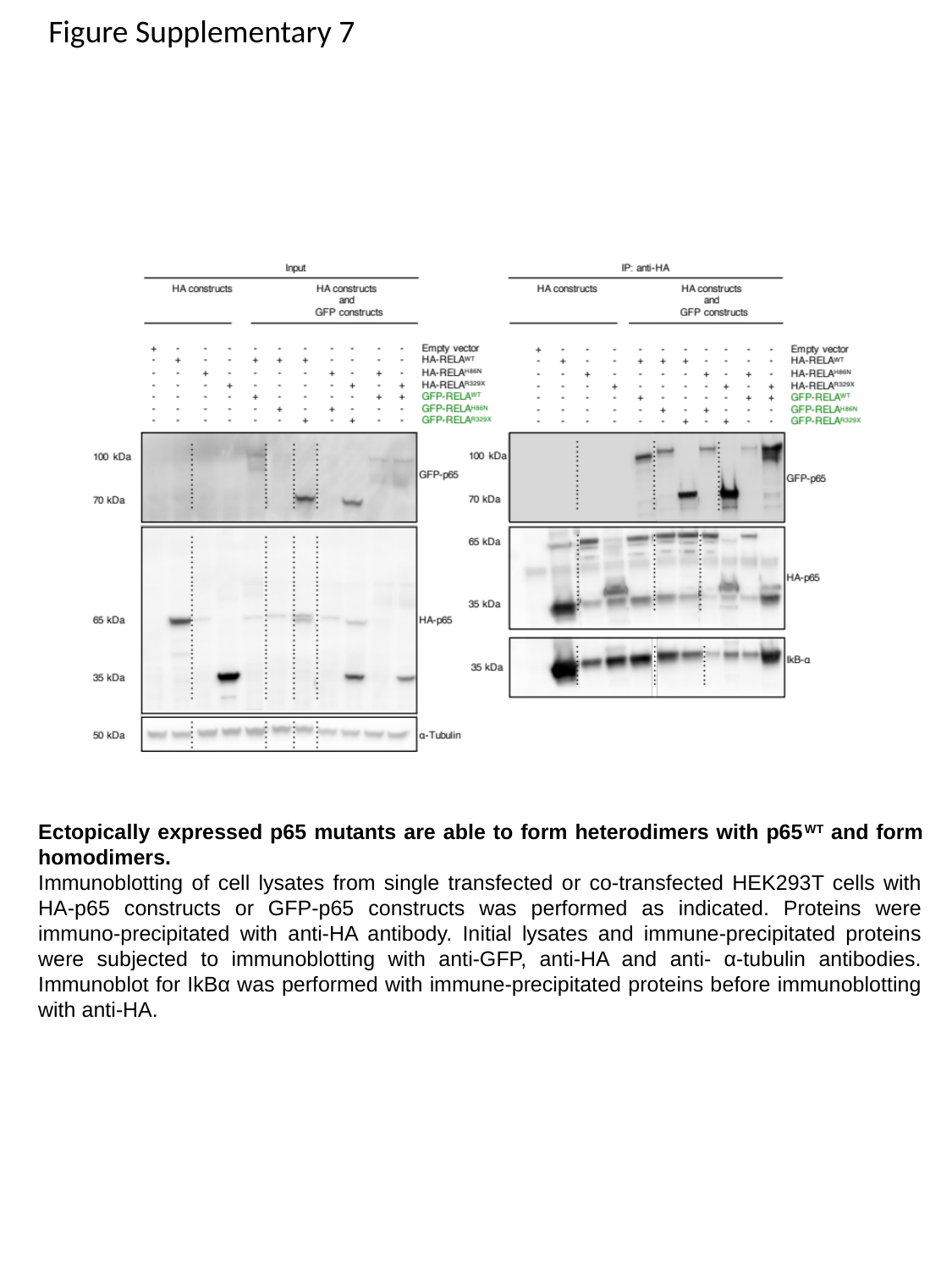

Figure Supplementary 7
Ectopically expressed p65 mutants are able to form heterodimers with p65^WT^ and form homodimers.
Immunoblotting of cell lysates from single transfected or co-transfected HEK293T cells with HA-p65 constructs or GFP-p65 constructs was performed as indicated. Proteins were immuno-precipitated with anti-HA antibody. Initial lysates and immune-precipitated proteins were subjected to immunoblotting with anti-GFP, anti-HA and anti- α-tubulin antibodies. Immunoblot for IkBα was performed with immune-precipitated proteins before immunoblotting with anti-HA.
