## Supplementary figures and images for "Heterozygous RELA mutations cause early-onset systemic lupus erythematosus by hijacking the NF-κB pathway towards transcriptional activation of type-I Interferon genes"

### Fig. S6

#### Slide 1
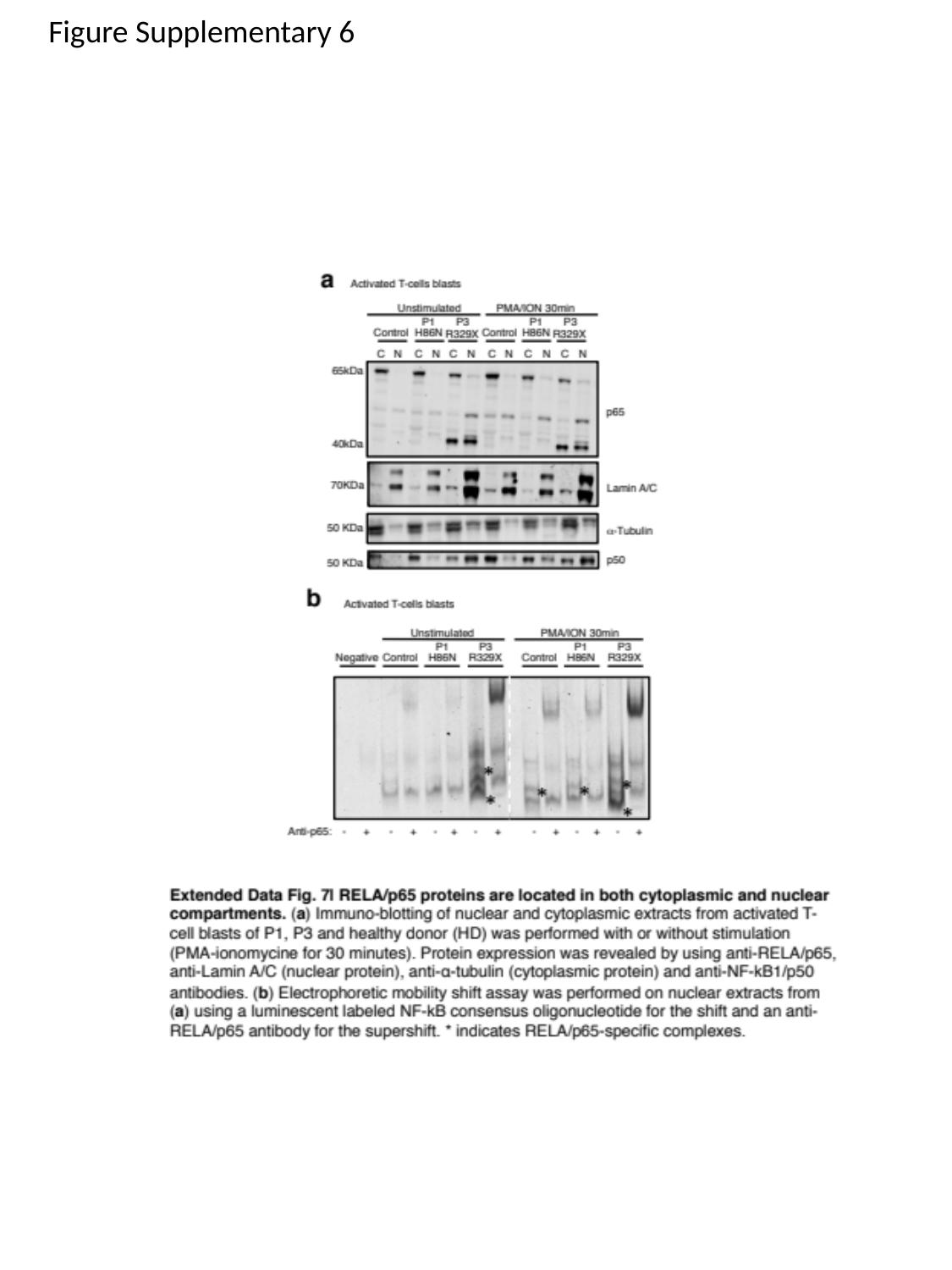

Figure Supplementary 6
