## Supplementary material for "Heterozygous RELA mutations cause early-onset systemic lupus erythematosus by hijacking the NF-κB pathway towards transcriptional activation of type-I Interferon genes": Fig. S5

### Slide 1
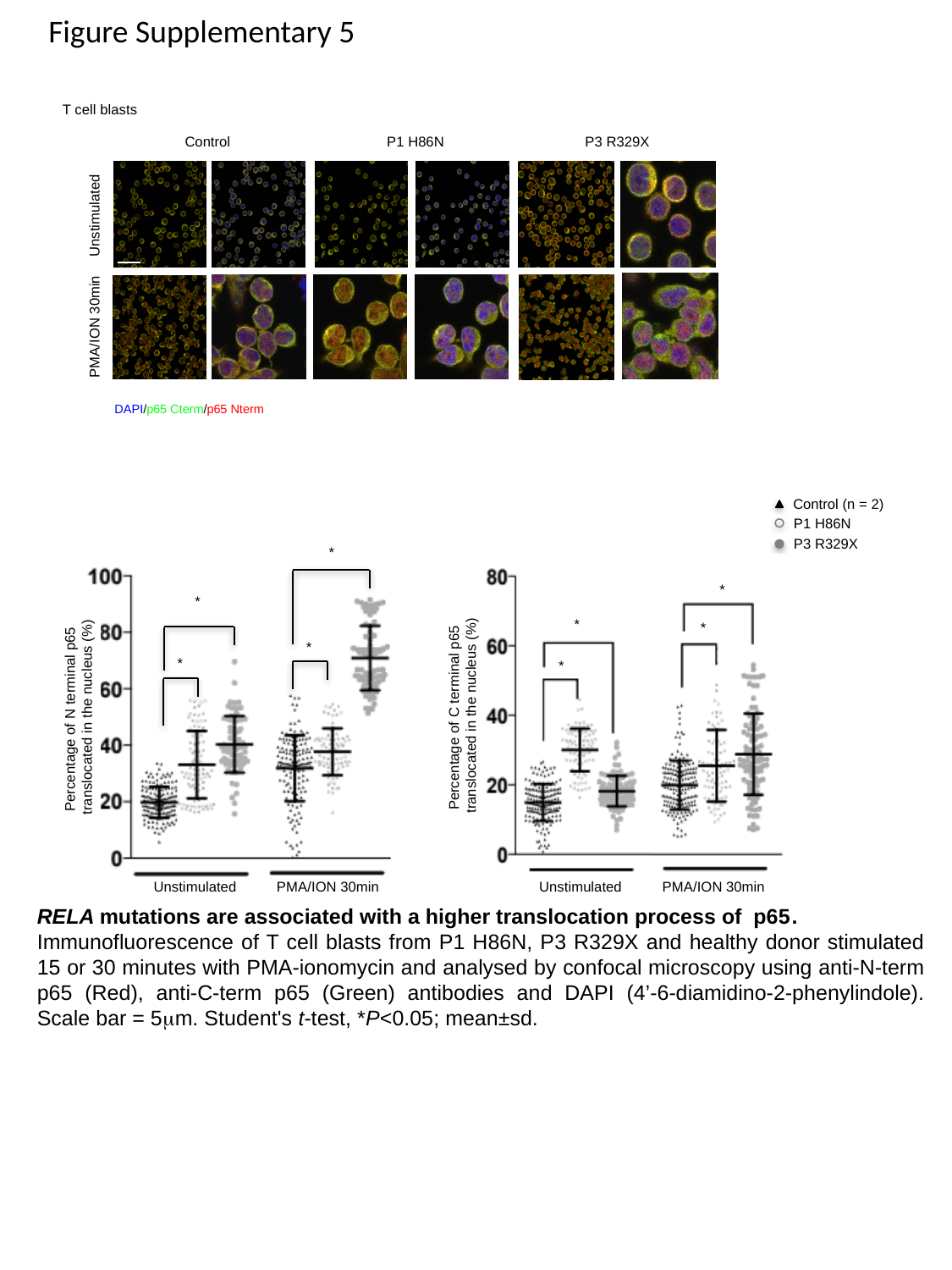

Figure Supplementary 5
T cell blasts
Control
P3 R329X
P1 H86N
Unstimulated
PMA/ION 30min
DAPI/p65 Cterm/p65 Nterm
Control (n = 2)
P1 H86N
P3 R329X
*
*
*
*
*
*
*
*
Percentage of C terminal p65
translocated in the nucleus (%)
Percentage of N terminal p65
translocated in the nucleus (%)
Unstimulated
Unstimulated
PMA/ION 30min
PMA/ION 30min
RELA mutations are associated with a higher translocation process of p65.
Immunofluorescence of T cell blasts from P1 H86N, P3 R329X and healthy donor stimulated 15 or 30 minutes with PMA-ionomycin and analysed by confocal microscopy using anti-N-term p65 (Red), anti-C-term p65 (Green) antibodies and DAPI (4’-6-diamidino-2-phenylindole). Scale bar = 5μm. Student's t-test, *P<0.05; mean±sd.
