## Supplementary material for "Heterozygous RELA mutations cause early-onset systemic lupus erythematosus by hijacking the NF-κB pathway towards transcriptional activation of type-I Interferon genes": Fig. S4

### Slide 1
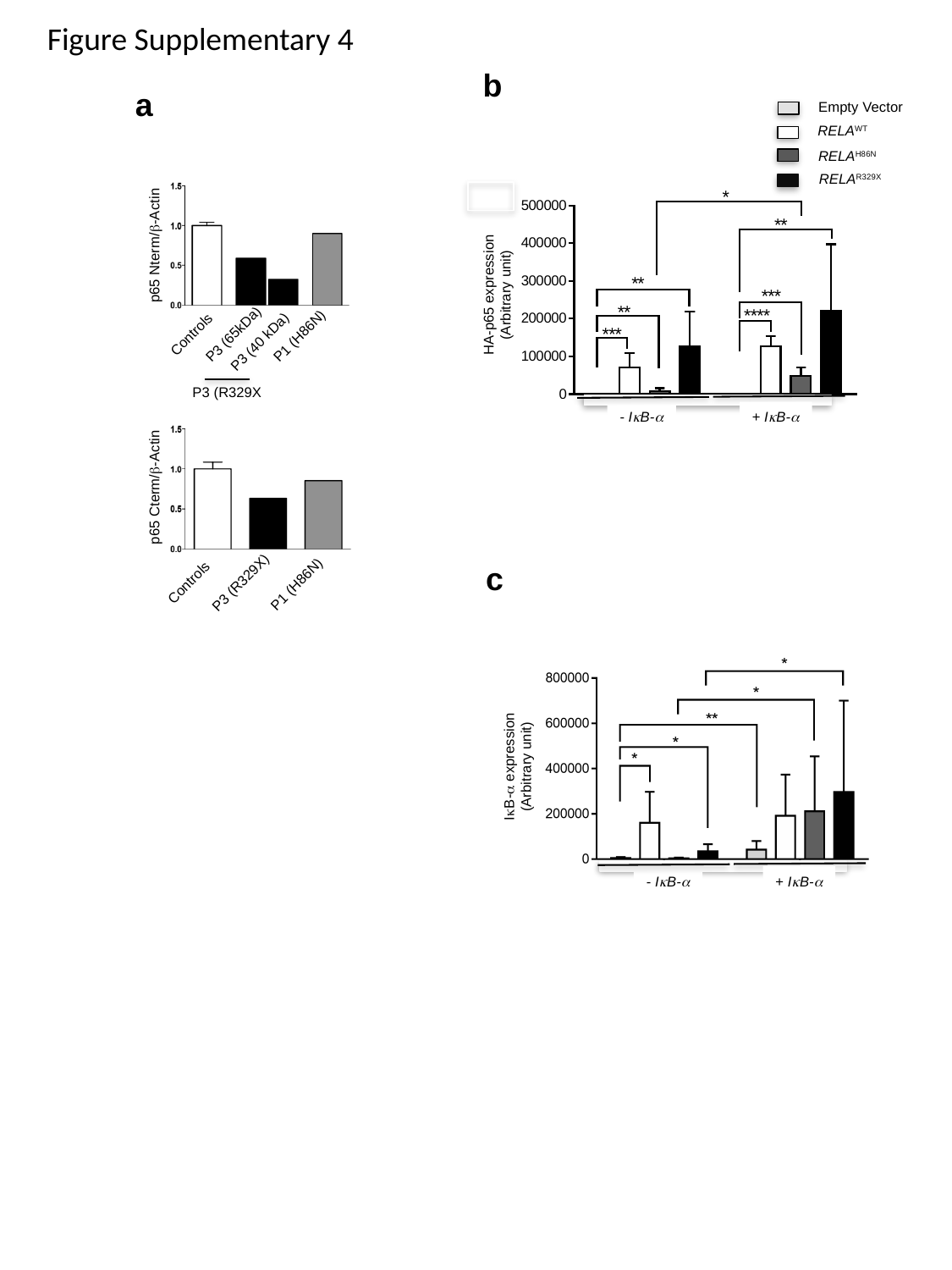

Figure Supplementary 4
b
Empty Vector
RELA^WT^
RELA^H86N^
RELA^R329X^
HA-p65 expression (Arbitrary unit)
- IκB-α
+ IκB-α
a
p65 Nterm/β-Actin
P3 (65kDa)
Controls
P1 (H86N)
P3 (40 kDa)
p65 Cterm/β-Actin
Controls
P3 (R329X)
P1 (H86N)
P3 (R329X
c
IκB-α expression (Arbitrary unit)
- IκB-α
+ IκB-α
