## Supplementary material for "Heterozygous RELA mutations cause early-onset systemic lupus erythematosus by hijacking the NF-κB pathway towards transcriptional activation of type-I Interferon genes": Fig S3

### Slide 1
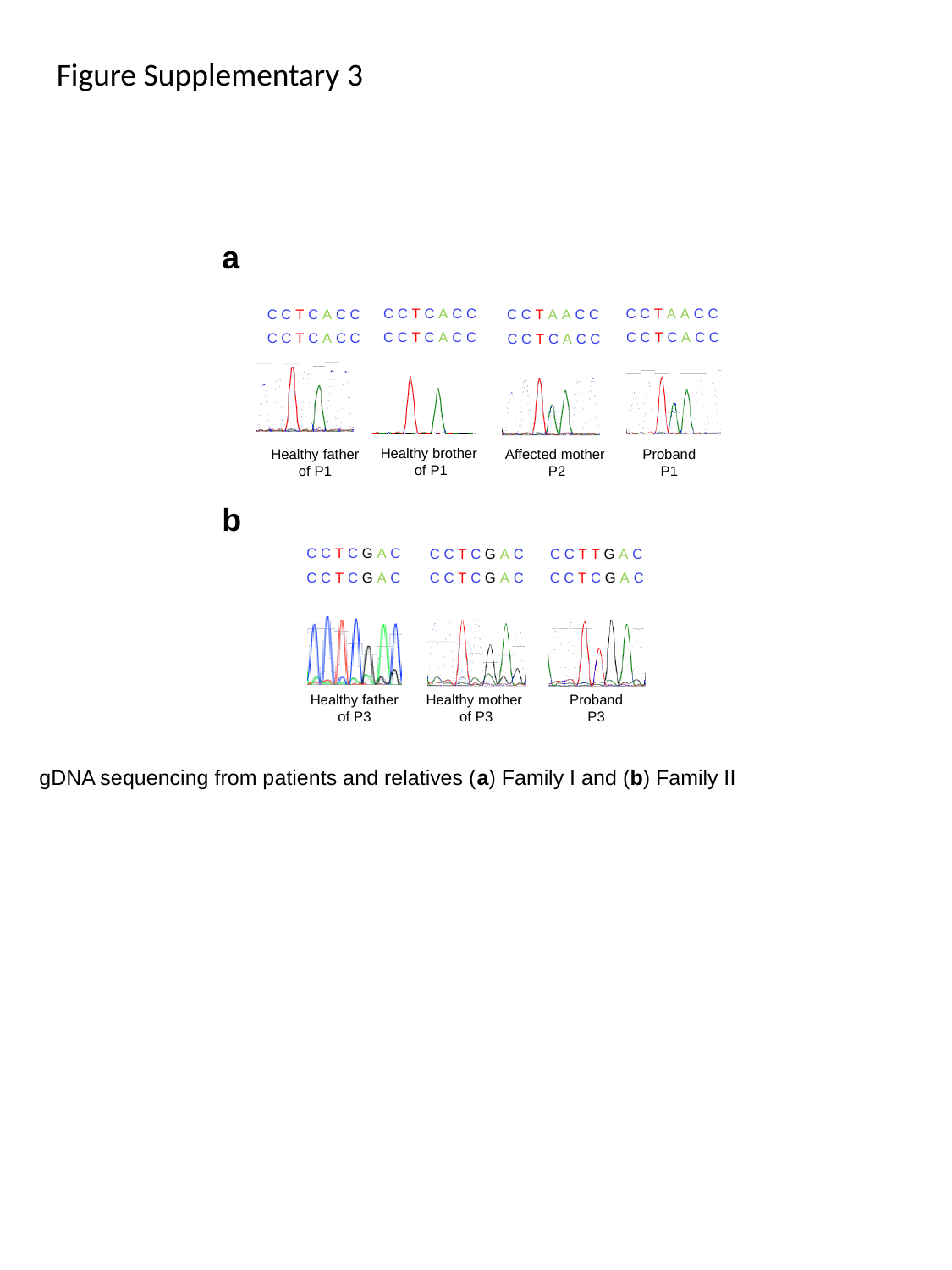

Figure Supplementary 3
a
C C T C A C C
C C T A A C C
C C T C A C C
C C T A A C C
C C T C A C C
C C T C A C C
C C T C A C C
C C T C A C C
Healthy brother
of P1
Proband
P1
Healthy father
of P1
Affected mother
P2
b
C C T C G A C
C C T C G A C
C C T T G A C
C C T C G A C
C C T C G A C
C C T C G A C
Healthy father
of P3
Healthy mother
of P3
Proband
P3
gDNA sequencing from patients and relatives (a) Family I and (b) Family II
