## Supplementary material for "Heterozygous RELA mutations cause early-onset systemic lupus erythematosus by hijacking the NF-κB pathway towards transcriptional activation of type-I Interferon genes": Fig. S2

### Slide 1
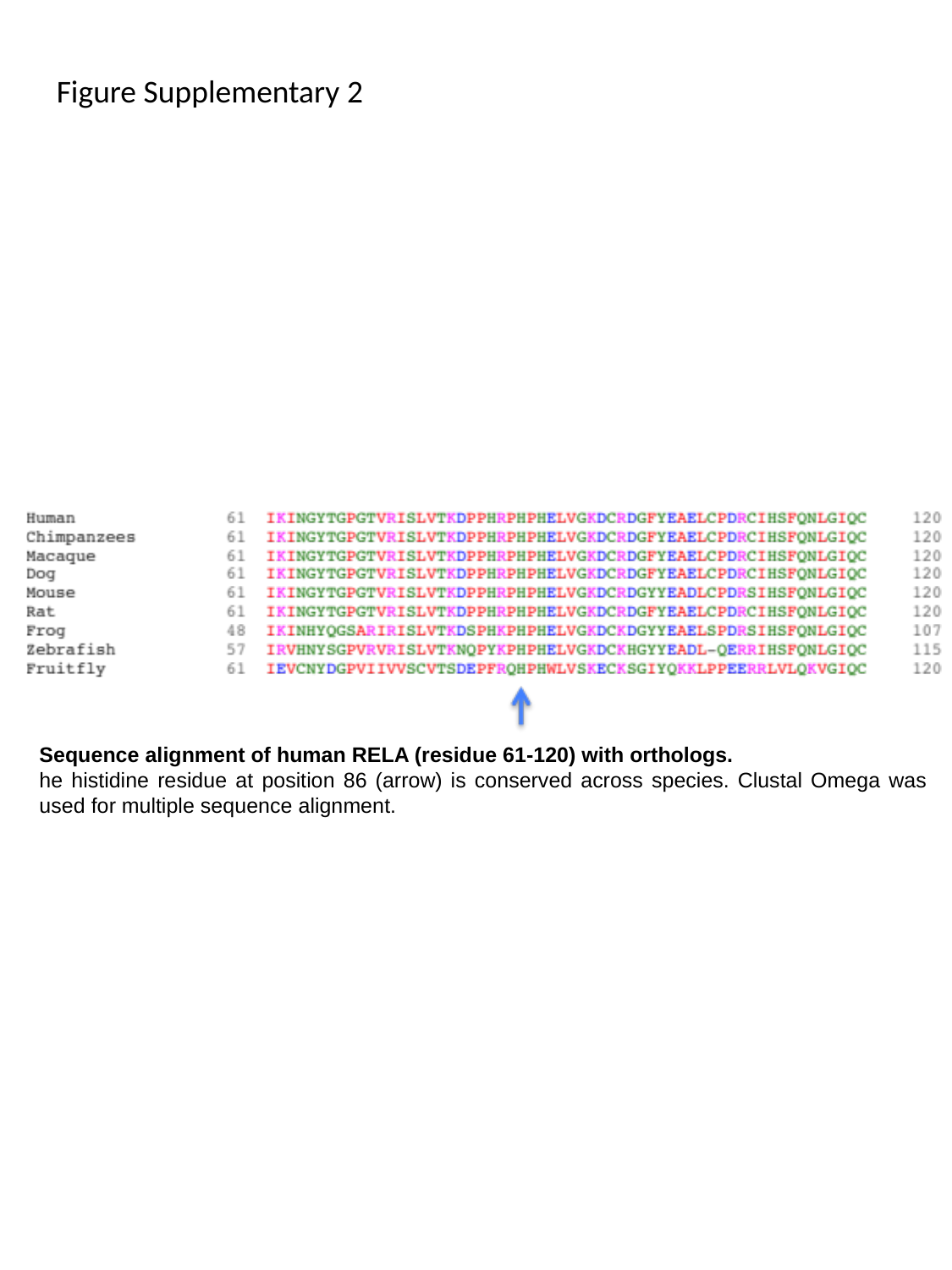

Figure Supplementary 2
Sequence alignment of human RELA (residue 61-120) with orthologs.
he histidine residue at position 86 (arrow) is conserved across species. Clustal Omega was used for multiple sequence alignment.
