## Supplementary material for "Heterozygous RELA mutations cause early-onset systemic lupus erythematosus by hijacking the NF-κB pathway towards transcriptional activation of type-I Interferon genes": Fig. S1

### Slide 1
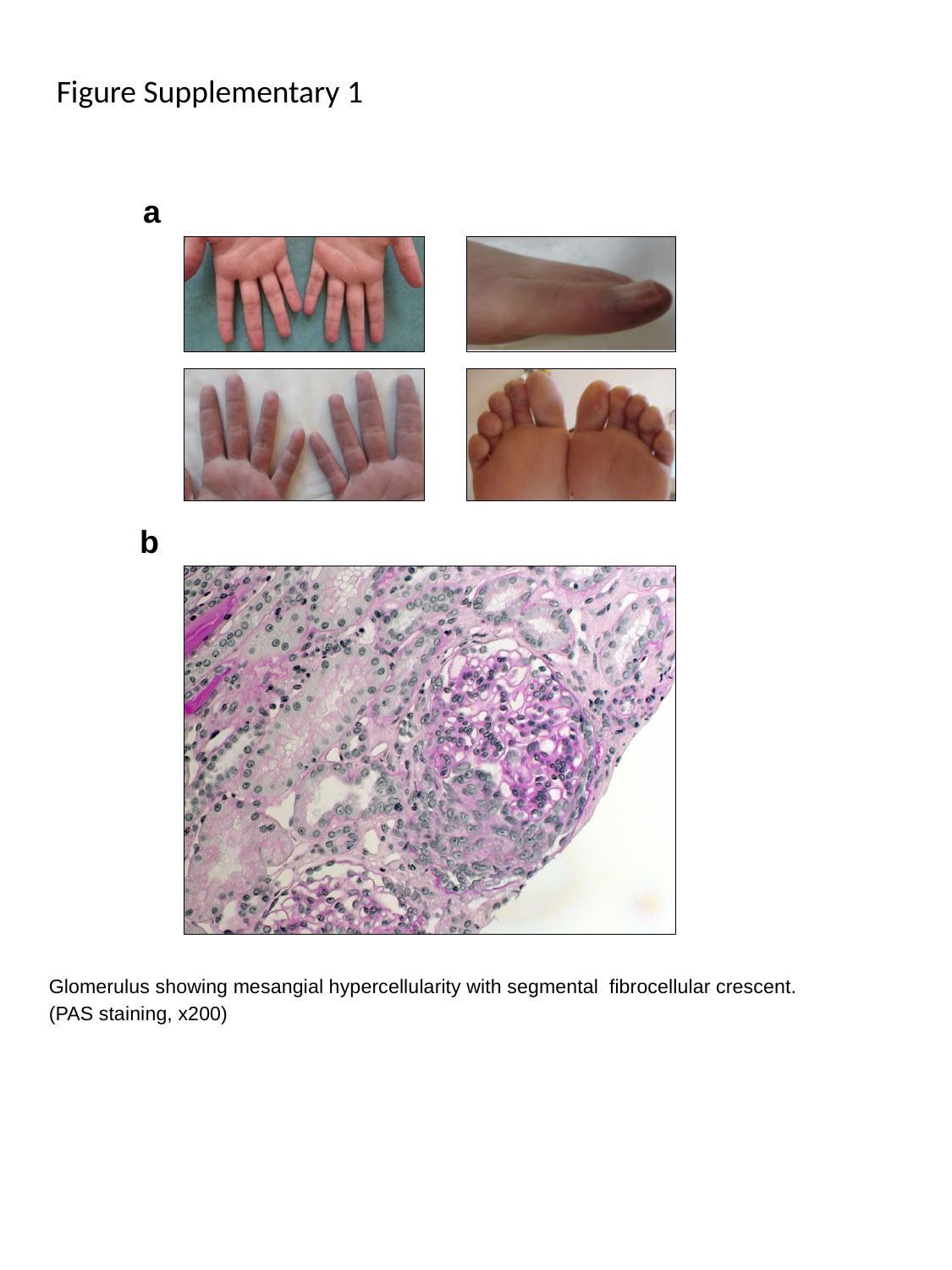

Figure Supplementary 1
a
b
Glomerulus showing mesangial hypercellularity with segmental fibrocellular crescent. (PAS staining, x200)
