## Supplementary material for "Heterozygous RELA mutations cause early-onset systemic lupus erythematosus by hijacking the NF-κB pathway towards transcriptional activation of type-I Interferon genes": Table S1

### Slide 1
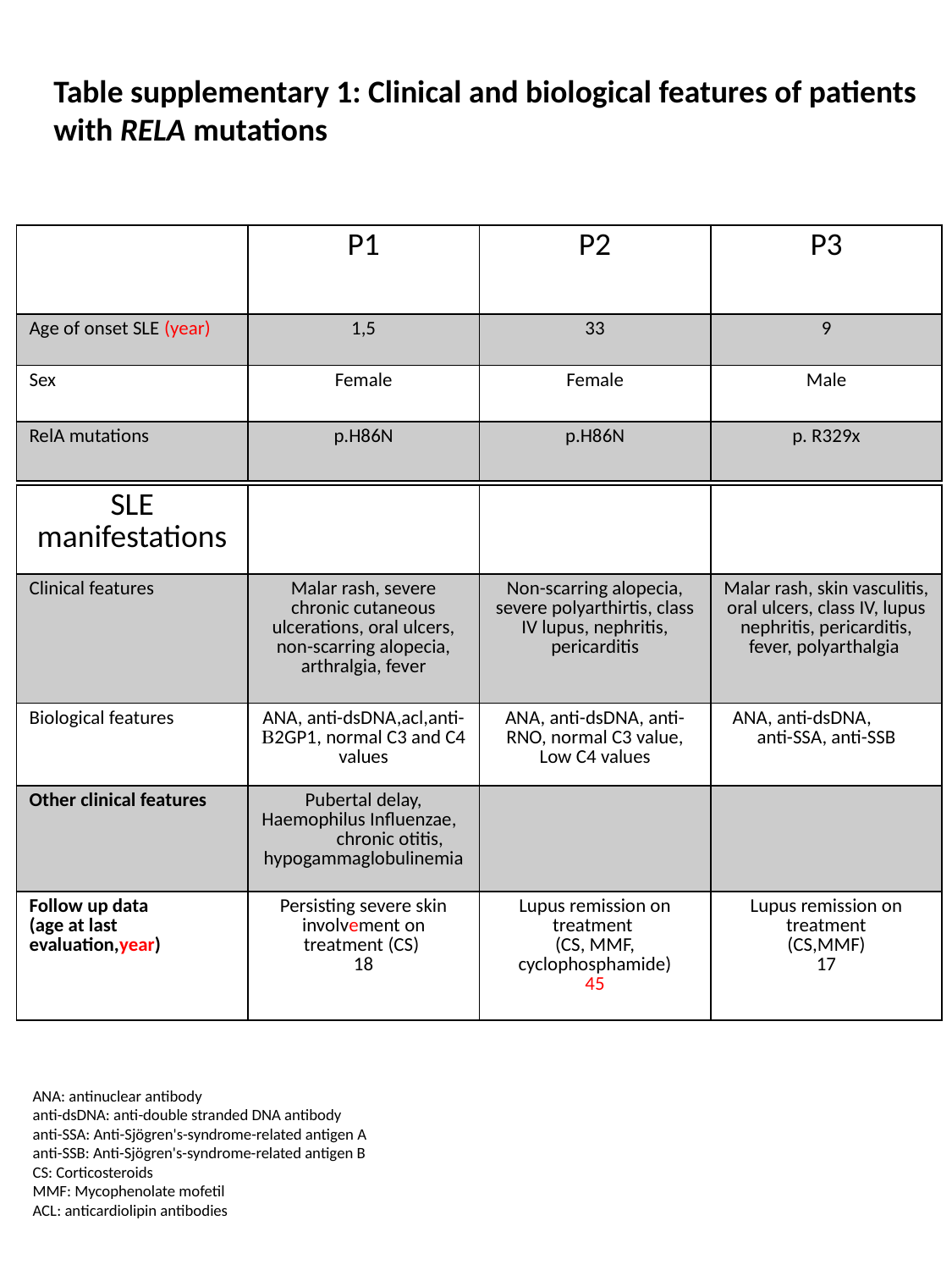

Table supplementary 1: Clinical and biological features of patients with RELA mutations
| | P1 | P2 | P3 |
| --- | --- | --- | --- |
| Age of onset SLE (year) | 1,5 | 33 | 9 |
| Sex | Female | Female | Male |
| RelA mutations | p.H86N | p.H86N | p. R329x |
| SLE manifestations | | | |
| --- | --- | --- | --- |
| Clinical features | Malar rash, severe chronic cutaneous ulcerations, oral ulcers, non-scarring alopecia, arthralgia, fever | Non-scarring alopecia, severe polyarthirtis, class IV lupus, nephritis, pericarditis | Malar rash, skin vasculitis, oral ulcers, class IV, lupus nephritis, pericarditis, fever, polyarthalgia |
| Biological features | ANA, anti-dsDNA,acl,anti-B2GP1, normal C3 and C4 values | ANA, anti-dsDNA, anti-RNO, normal C3 value, Low C4 values | ANA, anti-dsDNA, anti-SSA, anti-SSB |
| Other clinical features | Pubertal delay, Haemophilus Influenzae, chronic otitis, hypogammaglobulinemia | | |
| Follow up data (age at last evaluation,year) | Persisting severe skin involvement on treatment (CS) 18 | Lupus remission on treatment (CS, MMF, cyclophosphamide) 45 | Lupus remission on treatment (CS,MMF) 17 |
ANA: antinuclear antibody
anti-dsDNA: anti-double stranded DNA antibody
anti-SSA: Anti-Sjögren's-syndrome-related antigen A
anti-SSB: Anti-Sjögren's-syndrome-related antigen B
CS: Corticosteroids
MMF: Mycophenolate mofetil
ACL: anticardiolipin antibodies
