## Supplementary material for "Heterozygous RELA mutations cause early-onset systemic lupus erythematosus by hijacking the NF-κB pathway towards transcriptional activation of type-I Interferon genes": Table S2

### Slide 1
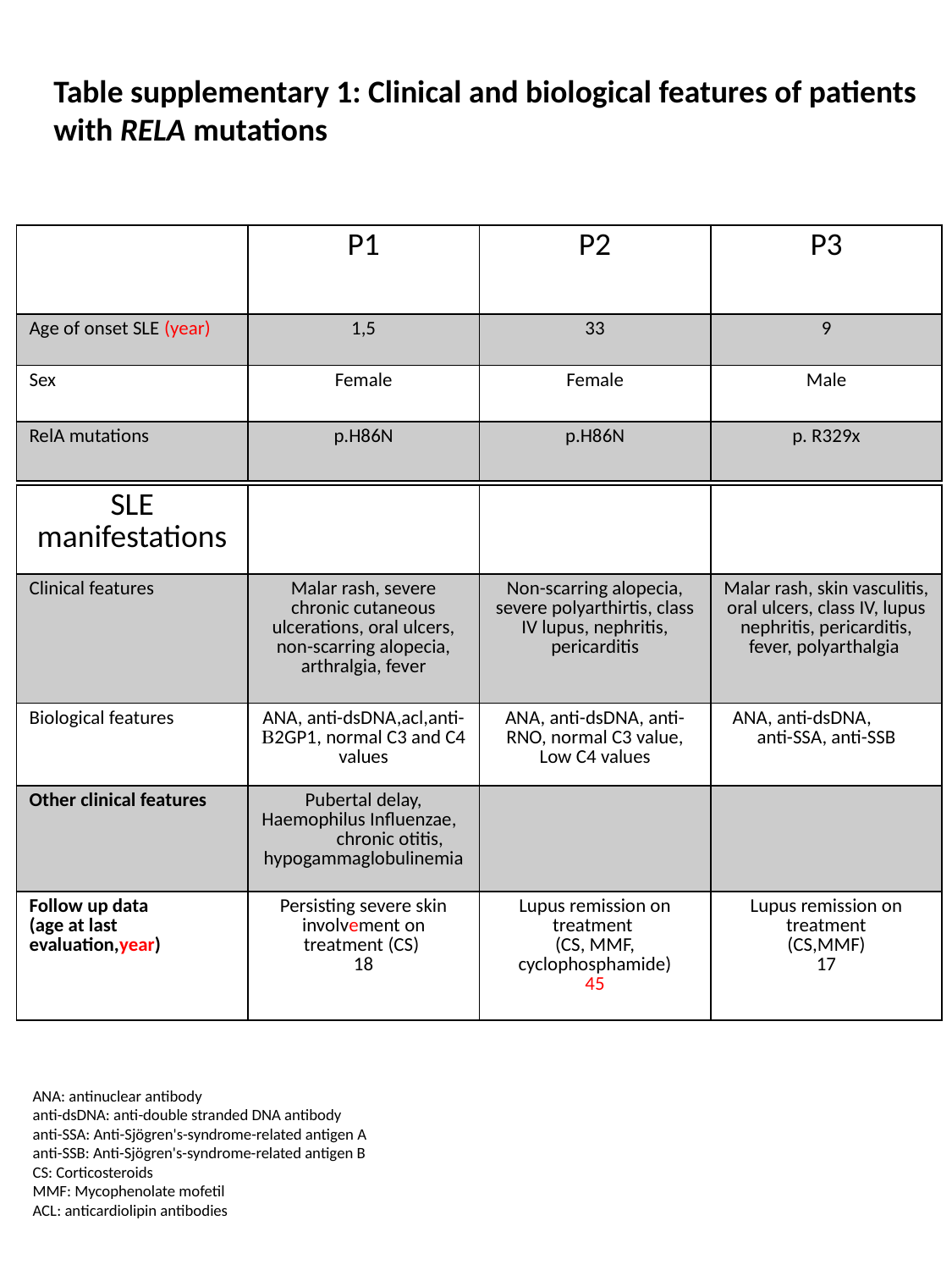

Table supplementary 1: Clinical and biological features of patients with RELA mutations
| | P1 | P2 | P3 |
| --- | --- | --- | --- |
| Age of onset SLE (year) | 1,5 | 33 | 9 |
| Sex | Female | Female | Male |
| RelA mutations | p.H86N | p.H86N | p. R329x |
| SLE manifestations | | | |
| --- | --- | --- | --- |
| Clinical features | Malar rash, severe chronic cutaneous ulcerations, oral ulcers, non-scarring alopecia, arthralgia, fever | Non-scarring alopecia, severe polyarthirtis, class IV lupus, nephritis, pericarditis | Malar rash, skin vasculitis, oral ulcers, class IV, lupus nephritis, pericarditis, fever, polyarthalgia |
| Biological features | ANA, anti-dsDNA,acl,anti-B2GP1, normal C3 and C4 values | ANA, anti-dsDNA, anti-RNO, normal C3 value, Low C4 values | ANA, anti-dsDNA, anti-SSA, anti-SSB |
| Other clinical features | Pubertal delay, Haemophilus Influenzae, chronic otitis, hypogammaglobulinemia | | |
| Follow up data (age at last evaluation,year) | Persisting severe skin involvement on treatment (CS) 18 | Lupus remission on treatment (CS, MMF, cyclophosphamide) 45 | Lupus remission on treatment (CS,MMF) 17 |
ANA: antinuclear antibody
anti-dsDNA: anti-double stranded DNA antibody
anti-SSA: Anti-Sjögren's-syndrome-related antigen A
anti-SSB: Anti-Sjögren's-syndrome-related antigen B
CS: Corticosteroids
MMF: Mycophenolate mofetil
ACL: anticardiolipin antibodies

### Slide 2
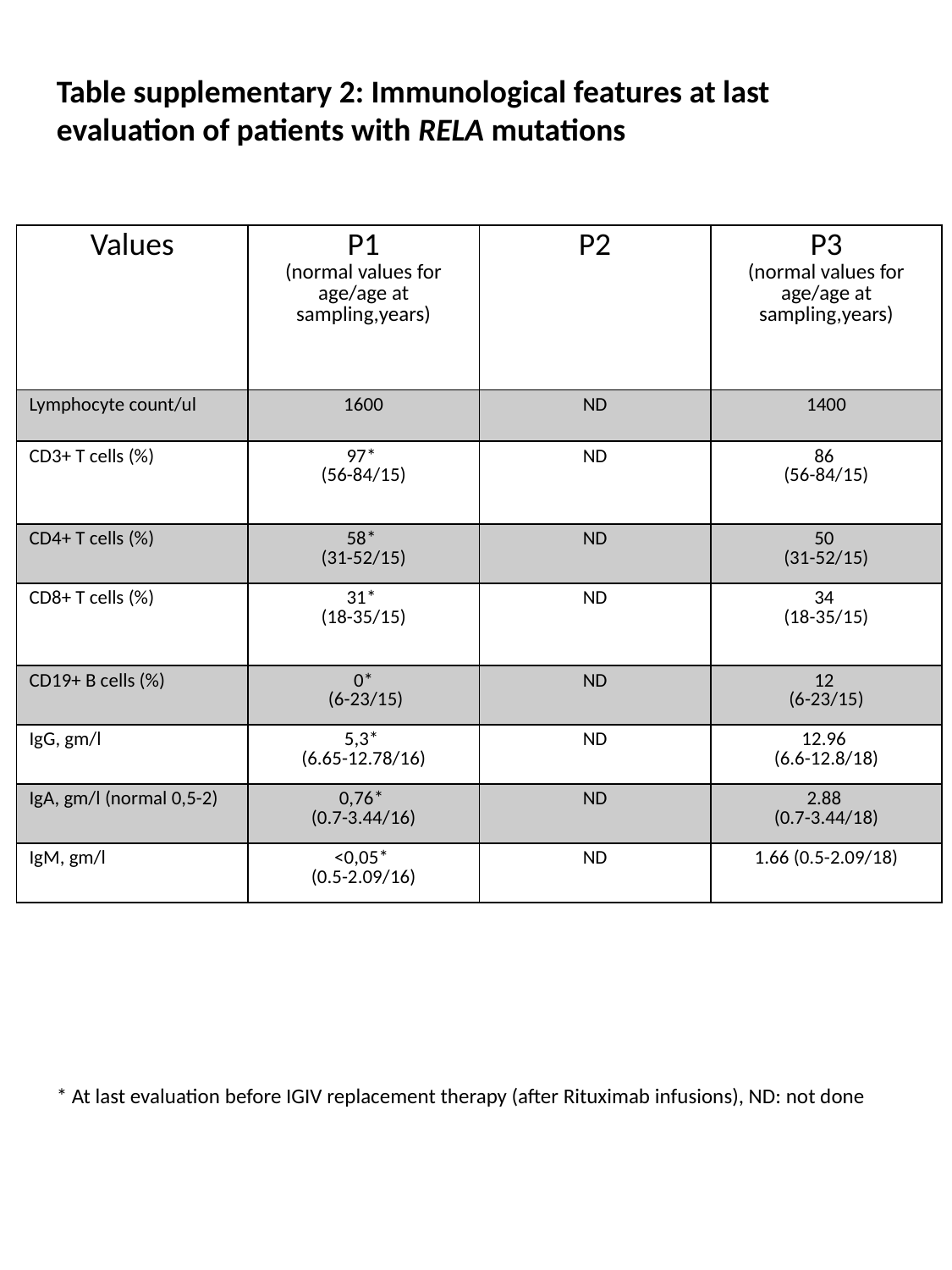

Table supplementary 2: Immunological features at last evaluation of patients with RELA mutations
| Values | P1 (normal values for age/age at sampling,years) | P2 | P3 (normal values for age/age at sampling,years) |
| --- | --- | --- | --- |
| Lymphocyte count/ul | 1600 | ND | 1400 |
| CD3+ T cells (%) | 97\* (56-84/15) | ND | 86 (56-84/15) |
| CD4+ T cells (%) | 58\* (31-52/15) | ND | 50 (31-52/15) |
| CD8+ T cells (%) | 31\* (18-35/15) | ND | 34 (18-35/15) |
| CD19+ B cells (%) | 0\* (6-23/15) | ND | 12 (6-23/15) |
| IgG, gm/l | 5,3\* (6.65-12.78/16) | ND | 12.96 (6.6-12.8/18) |
| IgA, gm/l (normal 0,5-2) | 0,76\* (0.7-3.44/16) | ND | 2.88 (0.7-3.44/18) |
| IgM, gm/l | <0,05\* (0.5-2.09/16) | ND | 1.66 (0.5-2.09/18) |
* At last evaluation before IGIV replacement therapy (after Rituximab infusions), ND: not done

### Slide 3
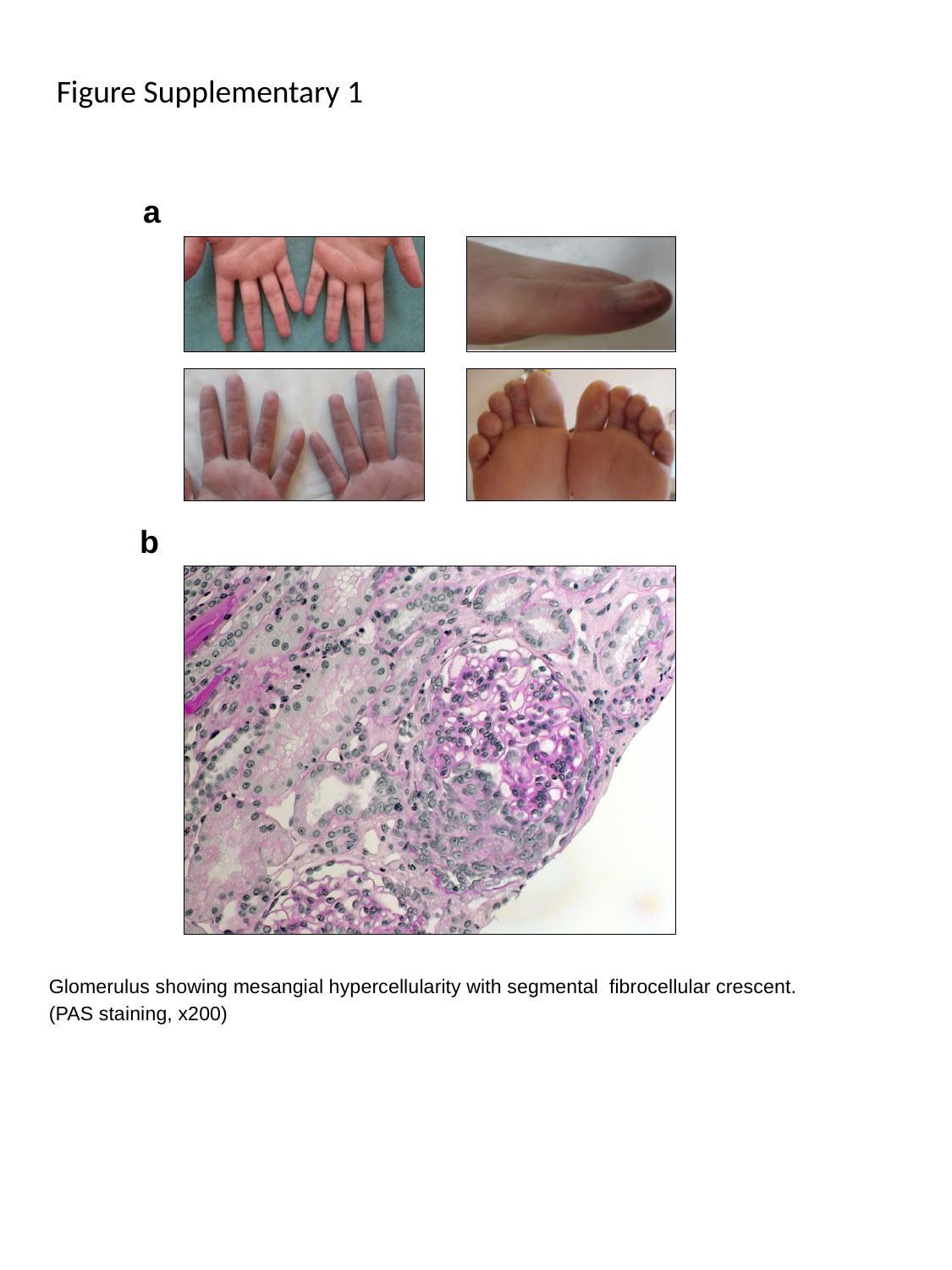

Figure Supplementary 1
a
b
Glomerulus showing mesangial hypercellularity with segmental fibrocellular crescent. (PAS staining, x200)

### Slide 4
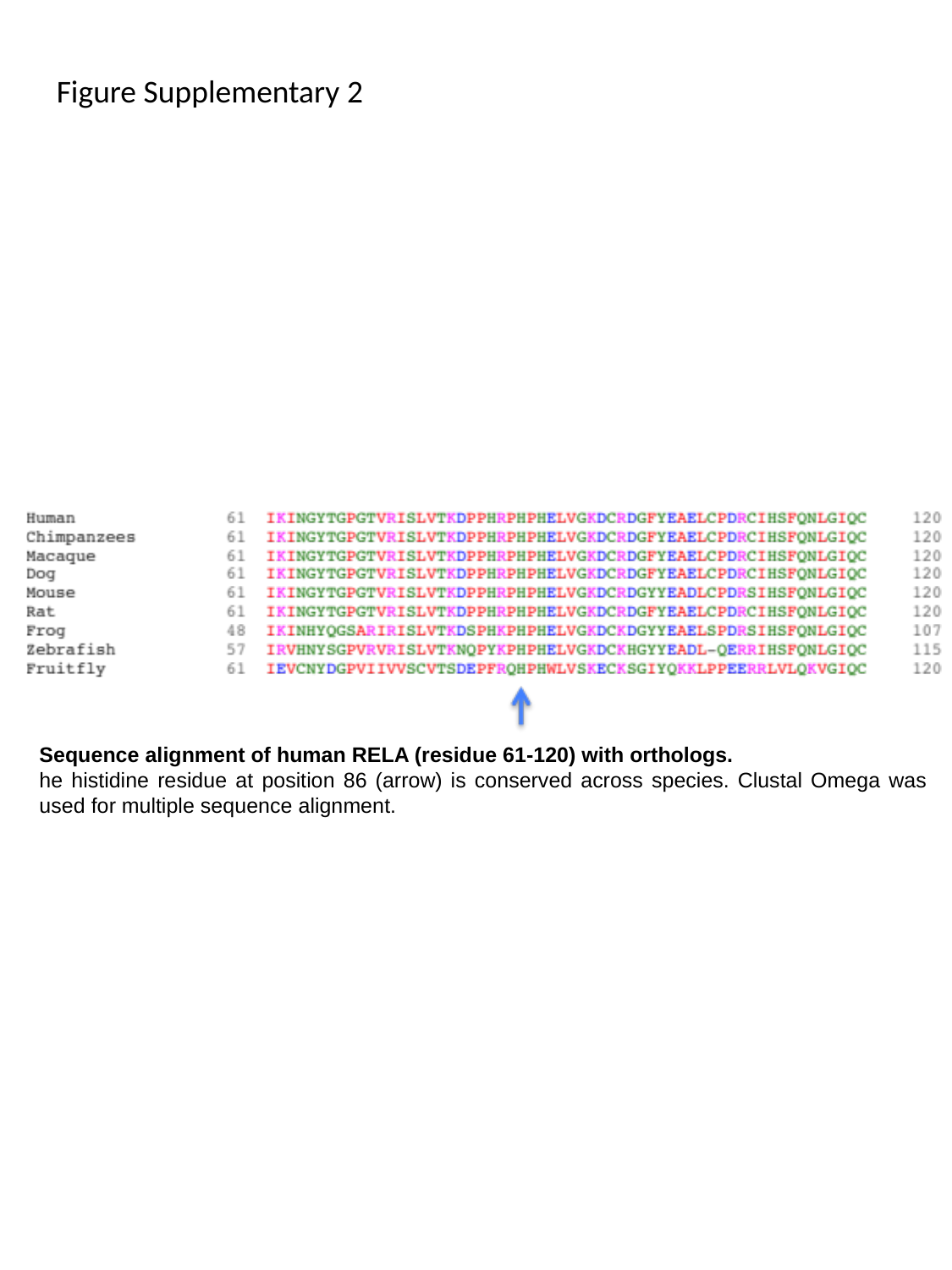

Figure Supplementary 2
Sequence alignment of human RELA (residue 61-120) with orthologs.
he histidine residue at position 86 (arrow) is conserved across species. Clustal Omega was used for multiple sequence alignment.

### Slide 5
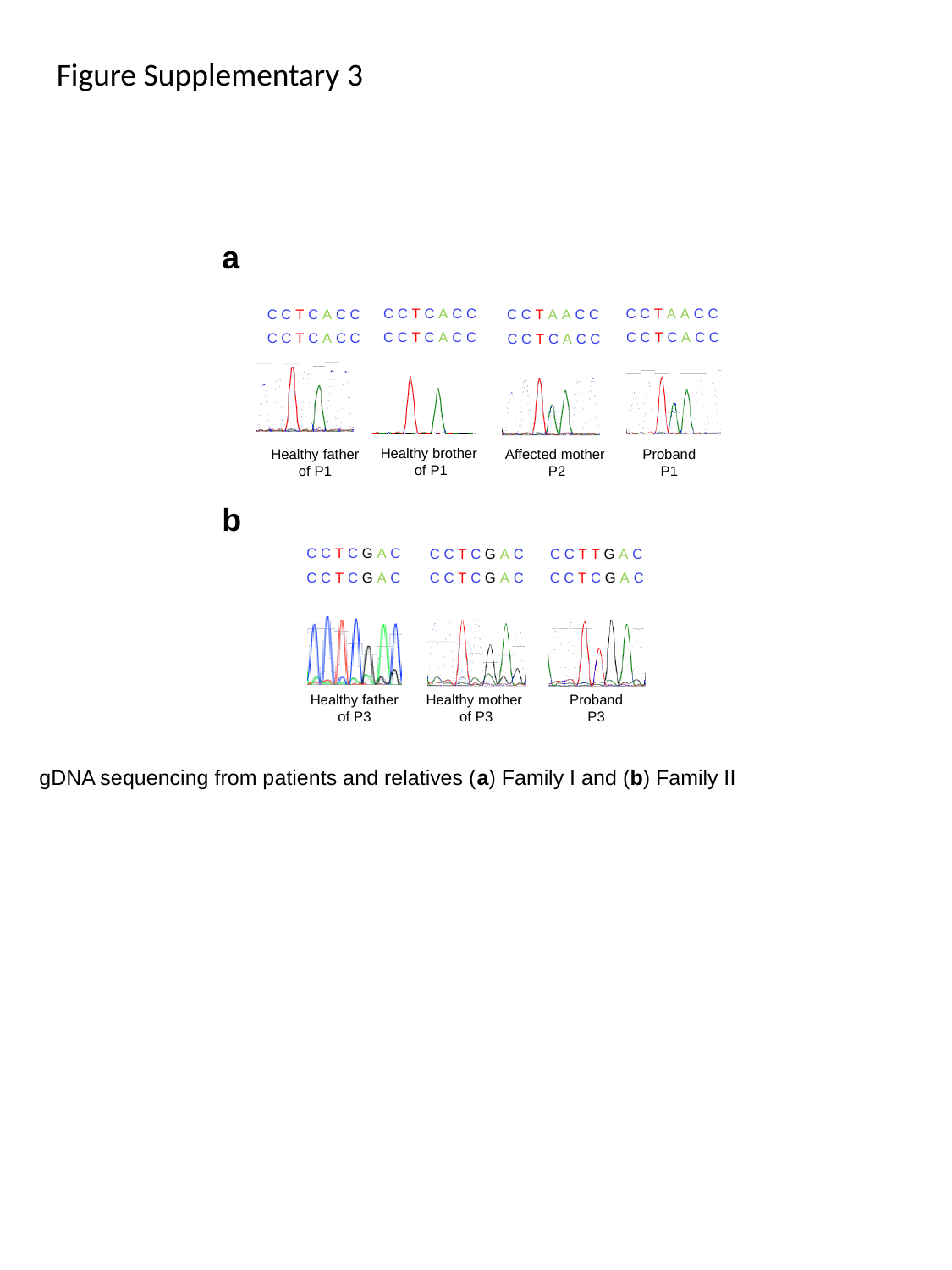

Figure Supplementary 3
a
C C T C A C C
C C T A A C C
C C T C A C C
C C T A A C C
C C T C A C C
C C T C A C C
C C T C A C C
C C T C A C C
Healthy brother
of P1
Proband
P1
Healthy father
of P1
Affected mother
P2
b
C C T C G A C
C C T C G A C
C C T T G A C
C C T C G A C
C C T C G A C
C C T C G A C
Healthy father
of P3
Healthy mother
of P3
Proband
P3
gDNA sequencing from patients and relatives (a) Family I and (b) Family II

### Slide 6
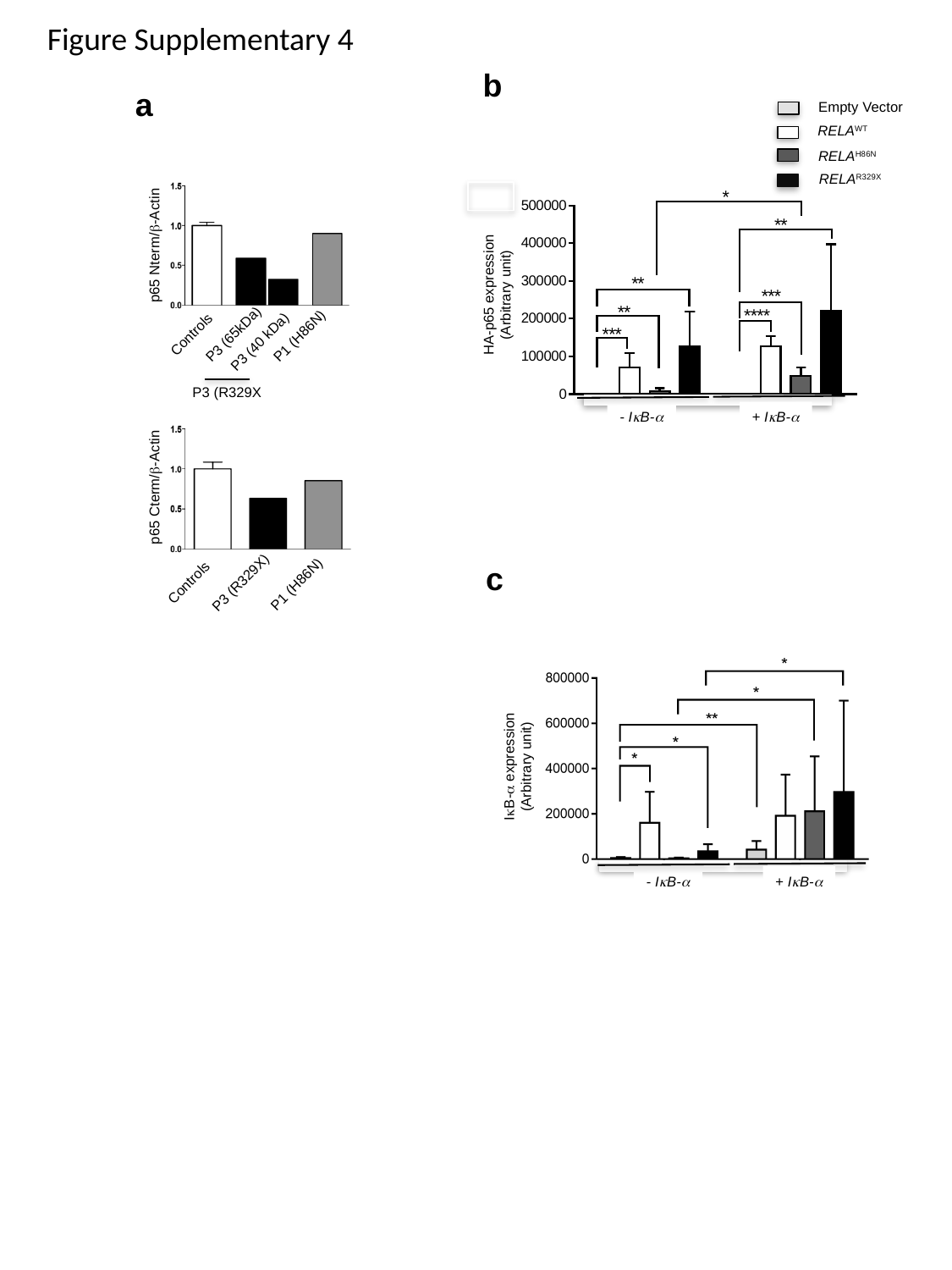

Figure Supplementary 4
b
Empty Vector
RELA^WT^
RELA^H86N^
RELA^R329X^
HA-p65 expression (Arbitrary unit)
- IκB-α
+ IκB-α
a
p65 Nterm/β-Actin
P3 (65kDa)
Controls
P1 (H86N)
P3 (40 kDa)
p65 Cterm/β-Actin
Controls
P3 (R329X)
P1 (H86N)
P3 (R329X
c
IκB-α expression (Arbitrary unit)
- IκB-α
+ IκB-α

### Slide 7
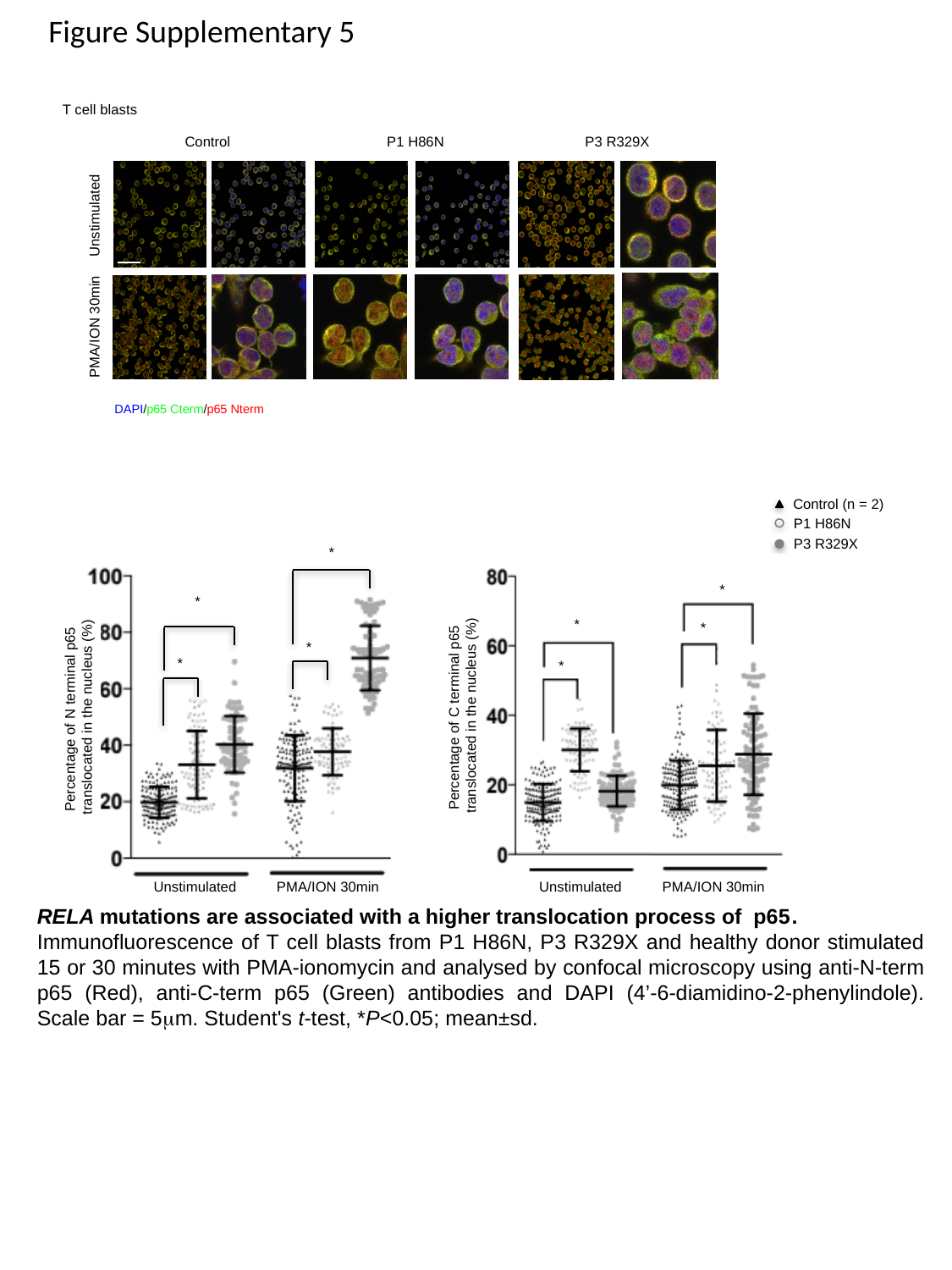

Figure Supplementary 5
T cell blasts
Control
P3 R329X
P1 H86N
Unstimulated
PMA/ION 30min
DAPI/p65 Cterm/p65 Nterm
Control (n = 2)
P1 H86N
P3 R329X
*
*
*
*
*
*
*
*
Percentage of C terminal p65
translocated in the nucleus (%)
Percentage of N terminal p65
translocated in the nucleus (%)
Unstimulated
Unstimulated
PMA/ION 30min
PMA/ION 30min
RELA mutations are associated with a higher translocation process of p65.
Immunofluorescence of T cell blasts from P1 H86N, P3 R329X and healthy donor stimulated 15 or 30 minutes with PMA-ionomycin and analysed by confocal microscopy using anti-N-term p65 (Red), anti-C-term p65 (Green) antibodies and DAPI (4’-6-diamidino-2-phenylindole). Scale bar = 5μm. Student's t-test, *P<0.05; mean±sd.

### Slide 8
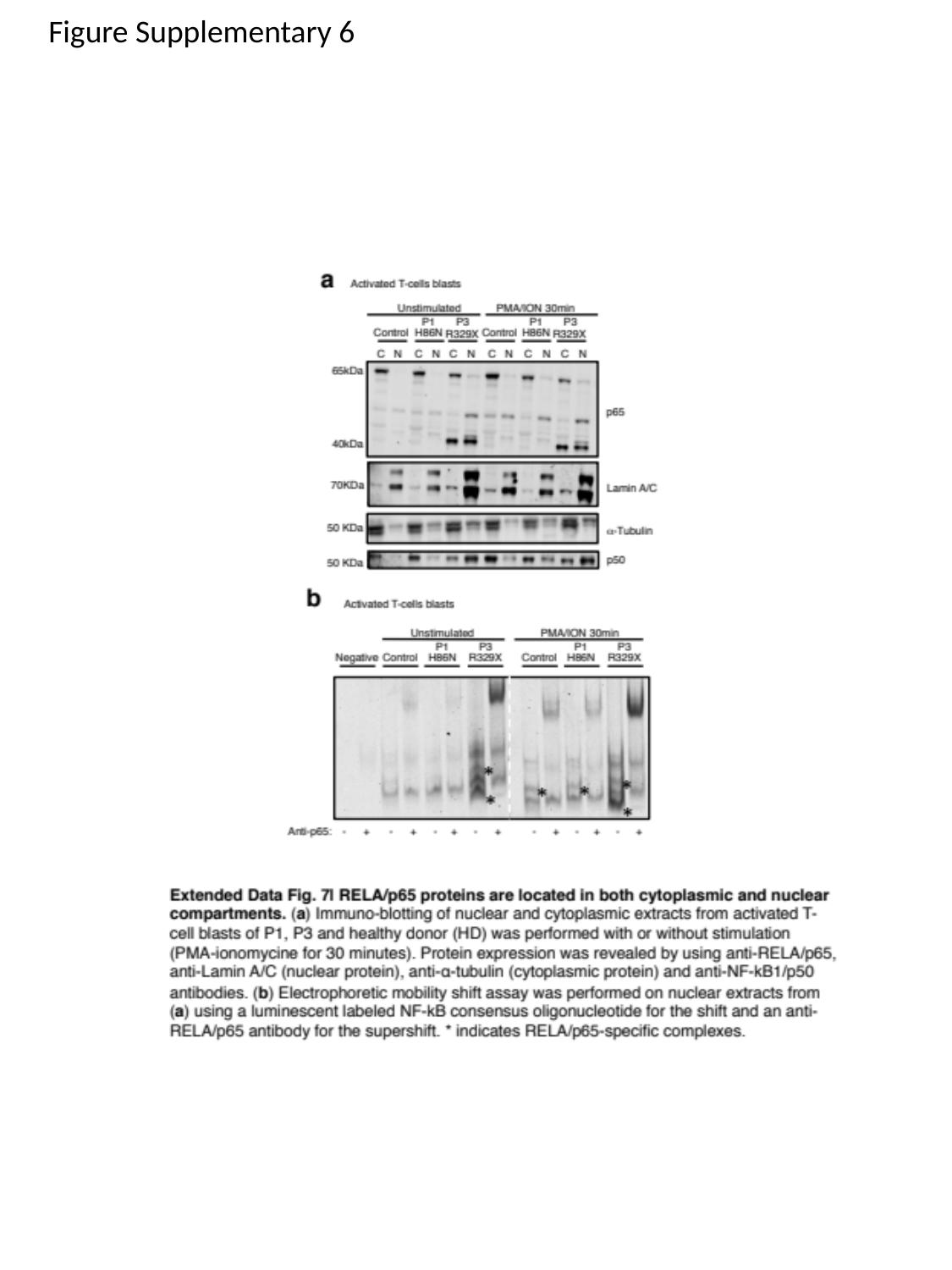

Figure Supplementary 6

### Slide 9
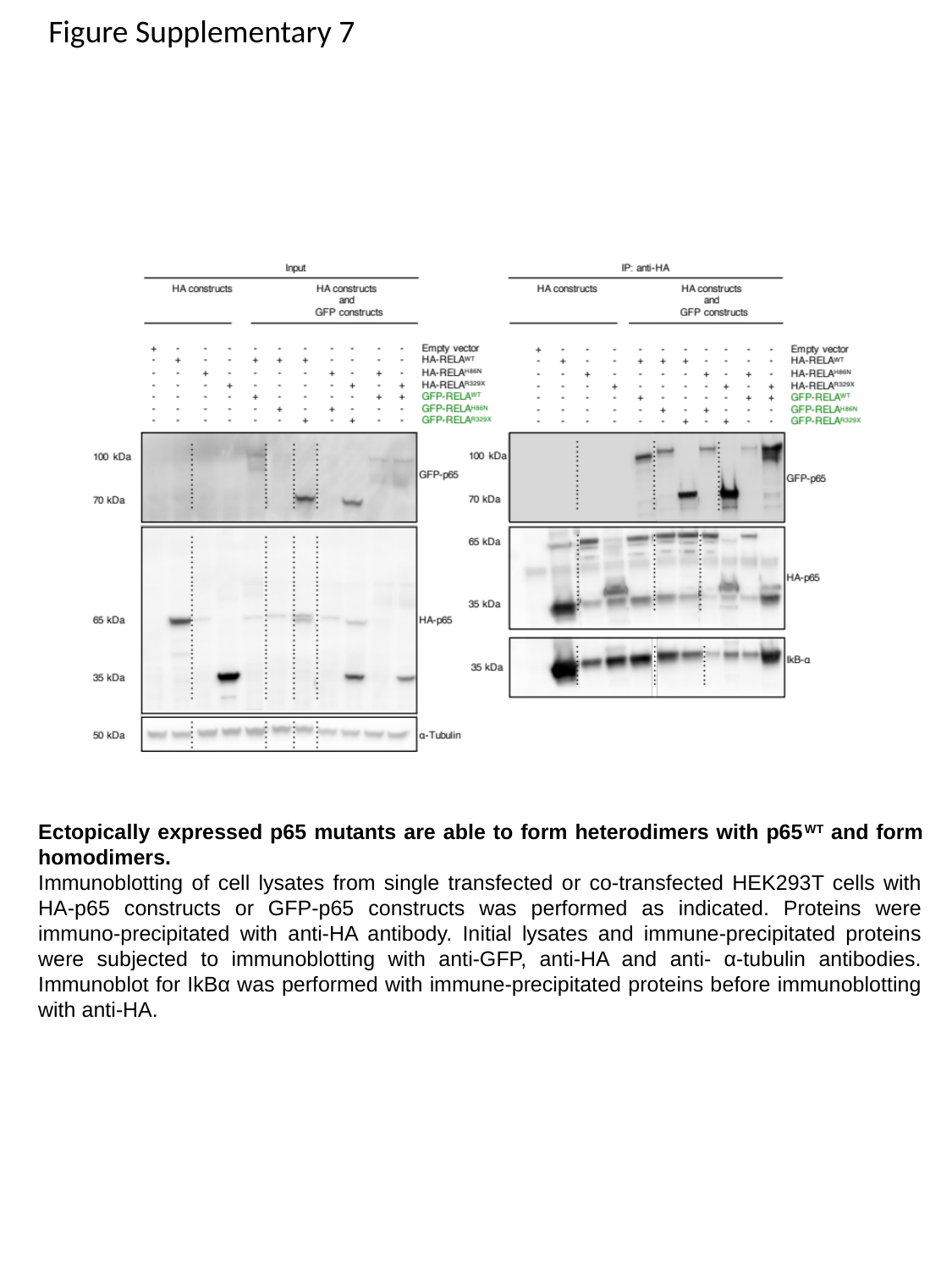

Figure Supplementary 7
Ectopically expressed p65 mutants are able to form heterodimers with p65^WT^ and form homodimers.
Immunoblotting of cell lysates from single transfected or co-transfected HEK293T cells with HA-p65 constructs or GFP-p65 constructs was performed as indicated. Proteins were immuno-precipitated with anti-HA antibody. Initial lysates and immune-precipitated proteins were subjected to immunoblotting with anti-GFP, anti-HA and anti- α-tubulin antibodies. Immunoblot for IkBα was performed with immune-precipitated proteins before immunoblotting with anti-HA.

### Slide 10
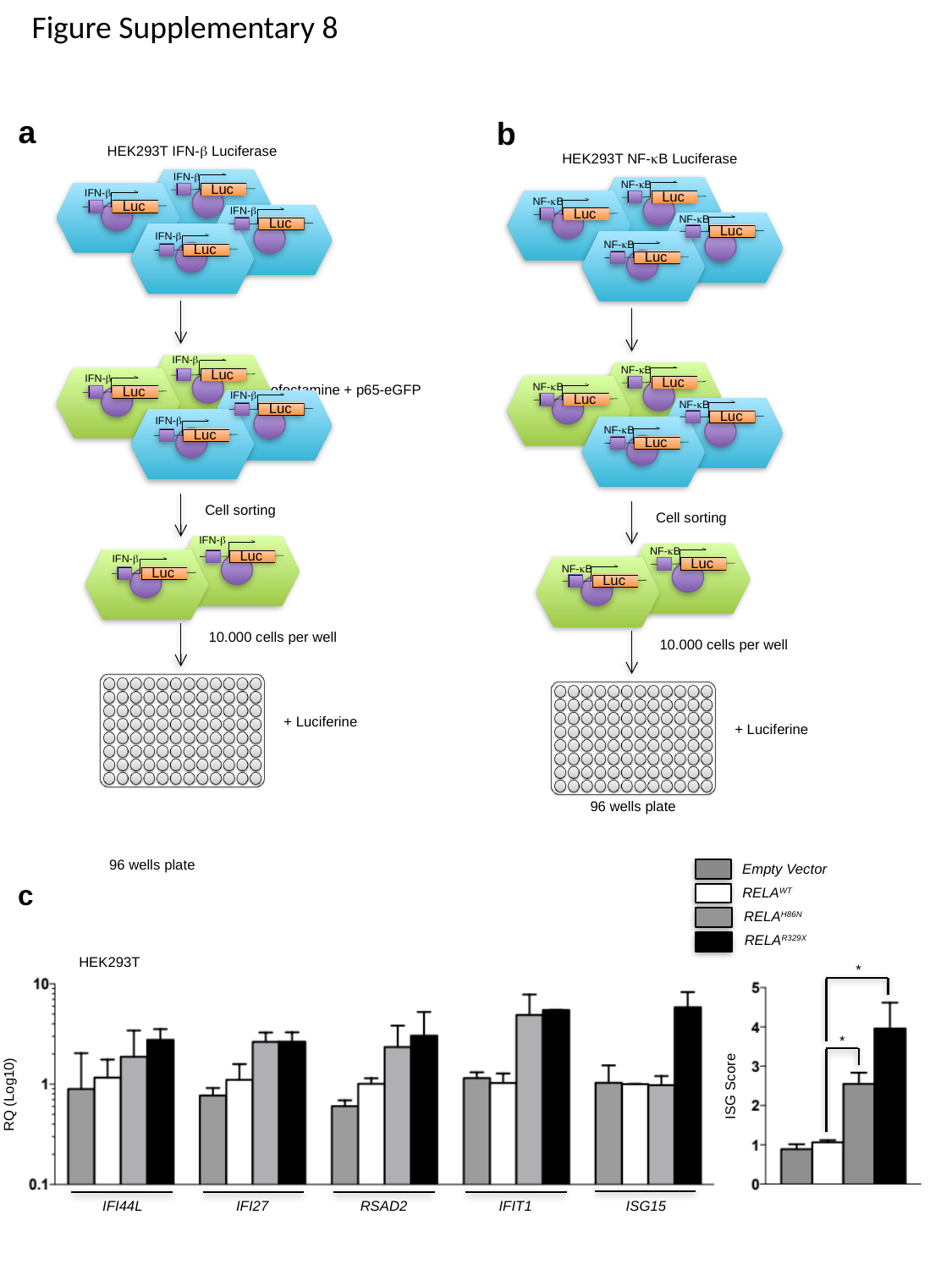

Figure Supplementary 8
a
b
HEK293T NF-κB Luciferase
NF-κB
Luc
NF-κB
Luc
NF-κB
Luc
NF-κB
Luc
NF-κB
Luc
NF-κB
Luc
NF-κB
Luc
NF-κB
Luc
Cell sorting
NF-κB
Luc
NF-κB
Luc
10.000 cells per well
+ Luciferine
96 wells plate
HEK293T IFN-β Luciferase
IFN-β
Luc
Luc
IFN-β
Luc
IFN-β
Luc
IFN-β
Luc
Luc
IFN-β
Luc
IFN-β
Luc
Cell sorting
IFN-β
Luc
IFN-β
Luc
IFN-β
IFN-β
10.000 cells per well
+ Luciferine
Lipofectamine + p65-eGFP
96 wells plate
Empty Vector
c
RELA^WT^
RELA^H86N^
RELA^R329X^
HEK293T
*
*
ISG Score
RQ (Log10)
IFI44L
IFI27
RSAD2
IFIT1
ISG15

### Slide 11
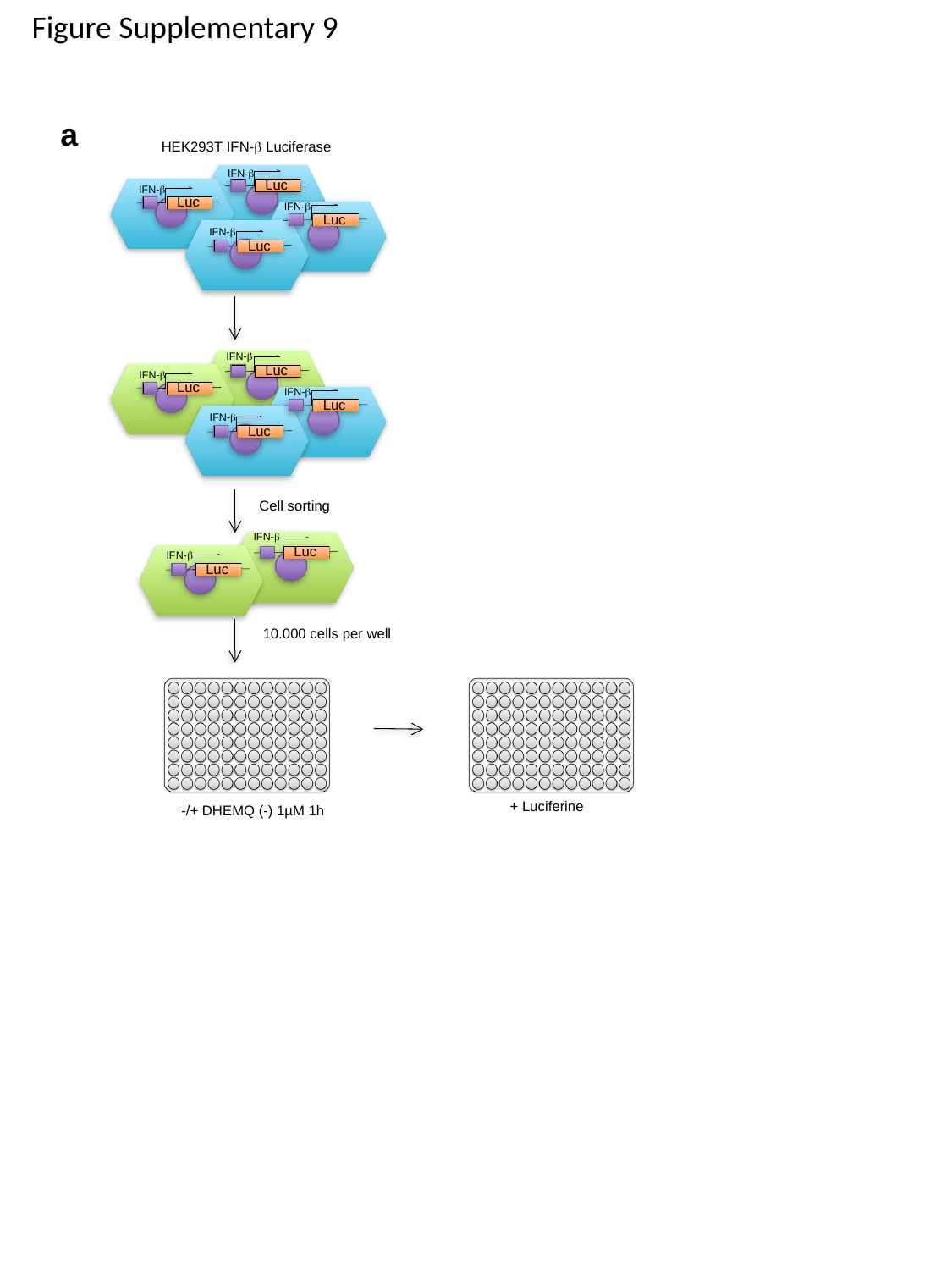

Figure Supplementary 9
a
HEK293T IFN-β Luciferase
IFN-β
Luc
Luc
IFN-β
Luc
IFN-β
Luc
IFN-β
Luc
Luc
IFN-β
Luc
IFN-β
Luc
Cell sorting
IFN-β
Luc
IFN-β
Luc
IFN-β
IFN-β
10.000 cells per well
+ Luciferine
-/+ DHEMQ (-) 1µM 1h

### Slide 12
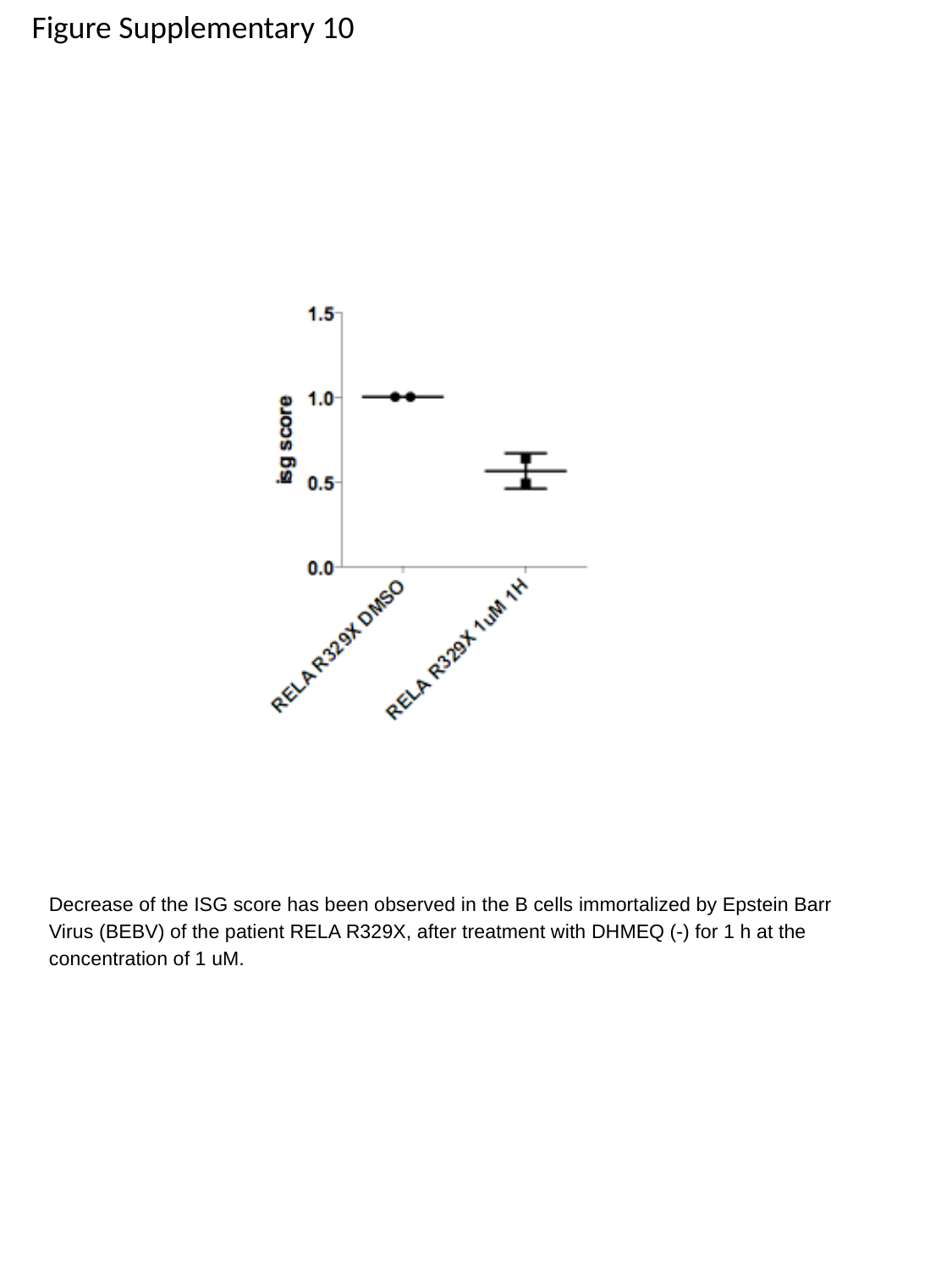

Figure Supplementary 10
Decrease of the ISG score has been observed in the B cells immortalized by Epstein Barr Virus (BEBV) of the patient RELA R329X, after treatment with DHMEQ (-) for 1 h at the concentration of 1 uM.
